# A multidimensional benchmark of confounder-aware differential abundance strategies for single- and multi-cohort microbiome studies

**DOI:** 10.64898/2026.10.29.762742

**Authors:** Xinyue Zhu, Lei Liu, Yiming Cao, Yunzhe Sheng, Bin Yang, Dingfeng Wu, Guoqing Zhang, Guoping Zhao, Na Jiao, Ruixin Zhu

**Affiliations:** Shanghai Key Laboratory of Maternal Fetal Medicine, Shanghai Institute of Maternal-Fetal Medicine and Gynecologic Oncology, Clinical and Translation Research Center, Shanghai First Maternity and Infant Hospital, School of Life Sciences and Technology, Tongji University, Shanghai 200092, P. R. China; State Key Laboratory of Genetic and Development of Complex Phenotypes, Fudan Microbiome Center, School of Life Sciences, Fudan University, Shanghai 200438, P. R. China; Shanghai Southgene Technology Co., Ltd., Shanghai 201203, P. R. China; National Genomics Data Center & Bio-Med Big Data Center, CAS Key Laboratory of Computational Biology, Shanghai Institute of Nutrition and Health, University of Chinese Academy of Sciences, Chinese Academy of Sciences, Shanghai 200031, P. R. China; School of Life Science, Hangzhou Institute for Advanced Study, University of Chinese Academy of Sciences, Hangzhou 310024, P. R. China

## Abstract

Confounding is a pervasive challenge in microbiome differential abundance analysis (DAA), yet systematic benchmarks of confounder-aware DAA remain lacking. Existing benchmarks have also largely evaluated statistical methods or analytical components in isolation rather than the complete analytical strategies used in practice. We evaluated 71 single-cohort strategies spanning default and extended input configurations using 2,500 real-data-informed simulations, reference-based recovery of established biological signals and resampling-based within-dataset consistency; benchmark-prioritized strategies were further assessed for stability under systematic perturbations. We further evaluated 63 multi-cohort extensions using simulations with cohort heterogeneity and empirical datasets. Covariate adjustment improved false-discovery control at the cost of reduced sensitivity, while performance also depended strongly on the specific combination of preprocessing procedure and statistical model. Different strategies were favored across benchmark dimensions and analytical objectives. ANCOM-BC2_CSS and edgeR_CSS were favored for high-confidence, perturbation-stable inference, whereas ANCOM-BC2_count and ANCOM-BC2_TMM provided more balanced performance. In multi-cohort settings, mega-analysis provided the strongest overall balance when individual-level data were available: ANCOM-BC2_CSS_Mega and edgeR_CSS_Mega showed stronger accuracy performance, whereas ANCOM-BC2_TMM_Mega and ANCOM-BC2_count_Mega showed greater cross-cohort consistency. We implemented these findings in xConDA, an interactive web server linking analytical objectives and study designs to benchmark-prioritized, confounder-adjusted microbiome DAA workflows.

## Introduction

Differential abundance analysis (DAA) is central to microbiome research and is widely used for identifying microbial features associated with host phenotypes, clinical states and environmental exposures. However, microbiome abundance data are strongly affected by biological and technical variables and by study-design factors, including inflammation, diet, medication use, sequencing depth, batch effects, and microbiome load [1–5]. When associated with both the phenotypes and microbial composition, these factors can generate statistically significant associations that do not represent phenotype-specific microbial signals [6, 7]. This issue is particularly evident in colorectal cancer (CRC), where several initially promising microbial biomarkers were later shown to be largely driven by inflammation, microbiome load, and other covariates rather than cancer status [3]. Related concerns have been reported across other microbiome-associated diseases [8–10]. Although matched designs, including comparisons with unaffected first-degree relatives, can partly reduce familial and shared environmental confounding [8], such samples are rarely available. Computational approaches such as miMatch [11] offer an alternative way to reduce host-related imbalance. Nevertheless, reliable association inference ultimately depends on how confounders are handled within the downstream DAA workflow.

Existing DAA methods differ in their statistical assumptions and in how they accommodate microbiome data characteristics such as compositionality, zero inflation, and overdispersion, and a subset supports explicit multivariable covariate adjustment [12–15]. In practice, these methods are applied to abundance profiles generated through different normalization and transformation procedures, which can alter the representation and distributional properties of the input data and consequently affect differential-abundance inference. These interacting choices mean that practical DAA analyses are better viewed as complete analytical strategies comprising preprocessing, statistical modelling and covariate adjustment, rather than as isolated statistical methods. More broadly, this reflects a common challenge in computational analysis, where individual algorithms are often benchmarked separately despite being deployed as components of multi-step workflows. Previous benchmarks have characterized method-specific performance and sensitivity–FDR trade-offs [15–22], and recent work has directly examined confounder adjustment under realistic simulation settings [16]. However, to our knowledge, no systematic benchmark has comprehensively evaluated confounder-aware DAA at the level of complete analytical strategies across diverse study settings. Evidence for selecting such strategies according to different analytical objectives therefore remains limited.

This challenge is amplified in multi-cohort microbiome studies [23–26]. Combining cohorts can increase statistical power and support assessment of association generalizability, but also introduces between-cohort heterogeneity, batch effects and incomplete covariate harmonization. Meta-analysis combines cohort-specific association statistics [27], whereas mega-analysis [28] jointly models pooled individual-level data while accounting for cohort structure. Cross-cohort integration therefore constitutes an additional computational design choice that can interact with the underlying single-cohort DAA strategy. Consequently, multi-cohort DAA performance depends not only on the underlying single-cohort strategy but also on the integration framework. Systematic evidence for selecting among these combinations remains limited.

Here, we established a multidimensional, strategy-level benchmark of confounder-aware microbiome DAA across single- and multi-cohort settings. We systematically assessed 71 end-to-end single-cohort strategies using real-data-informed simulation, reference-based recovery of established biological signals, resampling-based within-dataset consistency and stability under systematic perturbations. We then extended benchmark-prioritized strategies to 63 multi-cohort workflows based on meta- and mega-analysis and assessed their simulation-based accuracy and empirical cross-cohort consistency. Rather than seeking a universally optimal method, our framework uses multidimensional performance profiles to support objective-guide selection of strategies according to analytical priorities, study design and data availability. We further translated these findings into xConDA (https://www.biosino.org/xconda/), an interactive web server for benchmark-informed selection and execution of confounder-adjusted DAA workflows.

## Results

### 1 A multidimensional strategy-level framework for benchmarking confounder-aware microbiome DAA

To benchmark DAA as a complete analytical workflow rather than as isolated preprocessing or modelling components [29], we first constructed a strategy-level framework for confounder-aware microbiome DAA across single- and multi-cohort study designs. Each single-cohort strategy was defined as a prespecified combination of an input preprocessing scheme and a DAA method, with covariate adjustment implemented within the framework supported by that method (Fig. 1a, Table 1 and Supplementary Table 1). Pairing six preprocessing procedures with 14 widely used DAA methods and excluding statistically incompatible or recurrently unstable combinations yielded 71 feasible strategies. These included default input configurations based on developer-recommended workflows, together with extended input configurations representing additional preprocessing–model combinations, which enabled assessment of the effects of preprocessing choices.

**Fig. 1.**
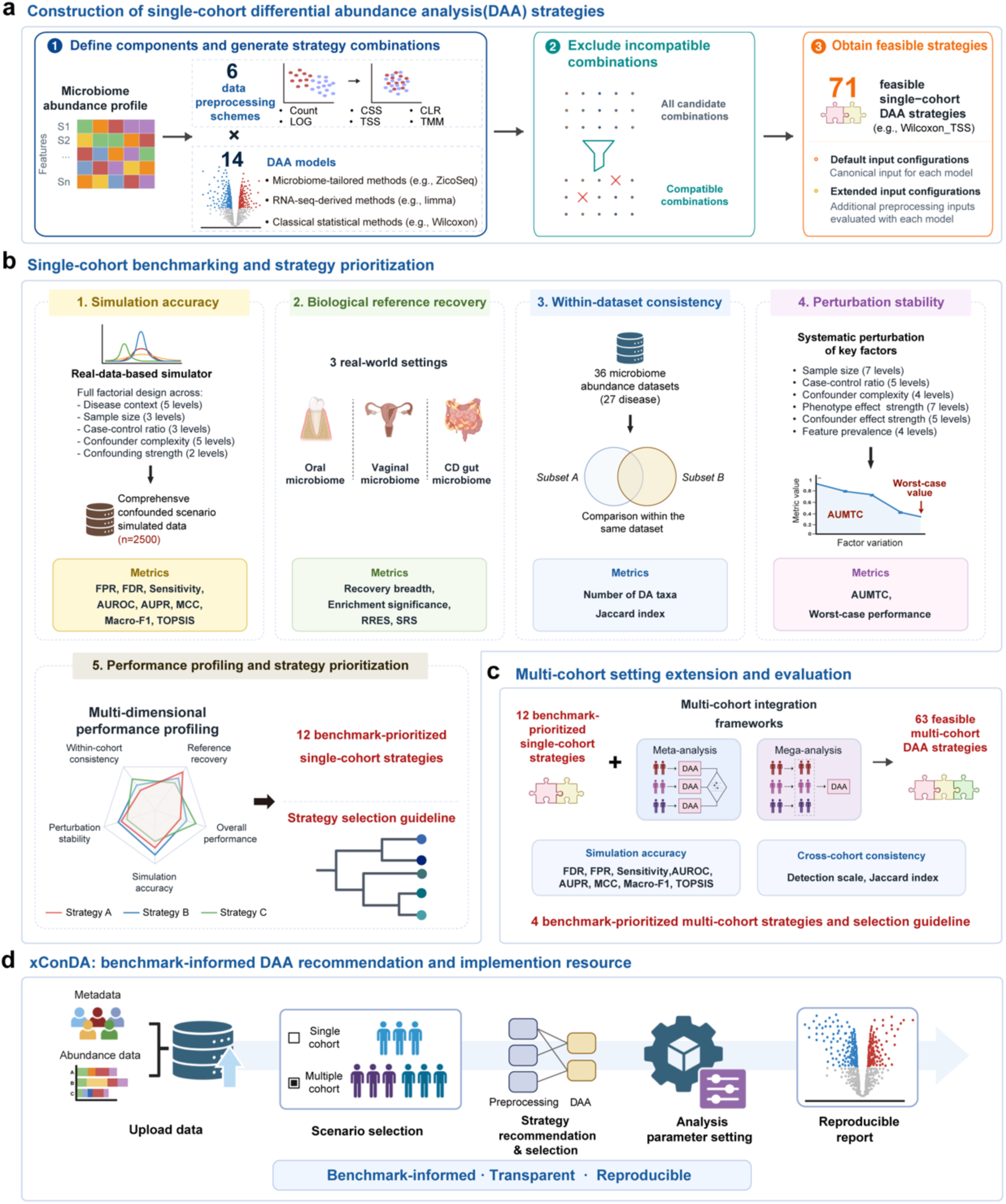
Study overview and multidimensional strategy-level benchmarking framework for confounder-aware DAA. **a**, Construction of single-cohort DAA strategies. Each strategy was defined as a combination of a data preprocessing procedure and a DAA statistical method. Across six preprocessing procedures and 14 DAA models, theoretically incompatible or computationally unstable combinations were excluded, yielding 71 executable strategies. **b**, Multidimensional benchmarking and prioritization of single-cohort strategies. Strategies were evaluated through a staged framework comprising broad assessment of simulation-based accuracy and reference-based recovery, followed by within-dataset consistency analysis and focused perturbation stress testing of prioritized strategies. Multidimensional performance profiles were used to prioritize strategies and to support strategy selection according to different analytical objectives. **c**, Extension to multi-cohort analysis. Benchmark-prioritized single-cohort strategies were combined with mega-analysis and meta-analysis frameworks to generate feasible multi-cohort DAA strategies, which were evaluated for simulation-based accuracy and cross-cohort consistency. **d**, xConDA implementation. xConDA translates the benchmark into an interactive web server for benchmark-informed selection and execution of confounder-adjusted DAA workflows, supporting user data upload, study and objective specification, executable analysis and reproducible reporting.

**Table 1.**
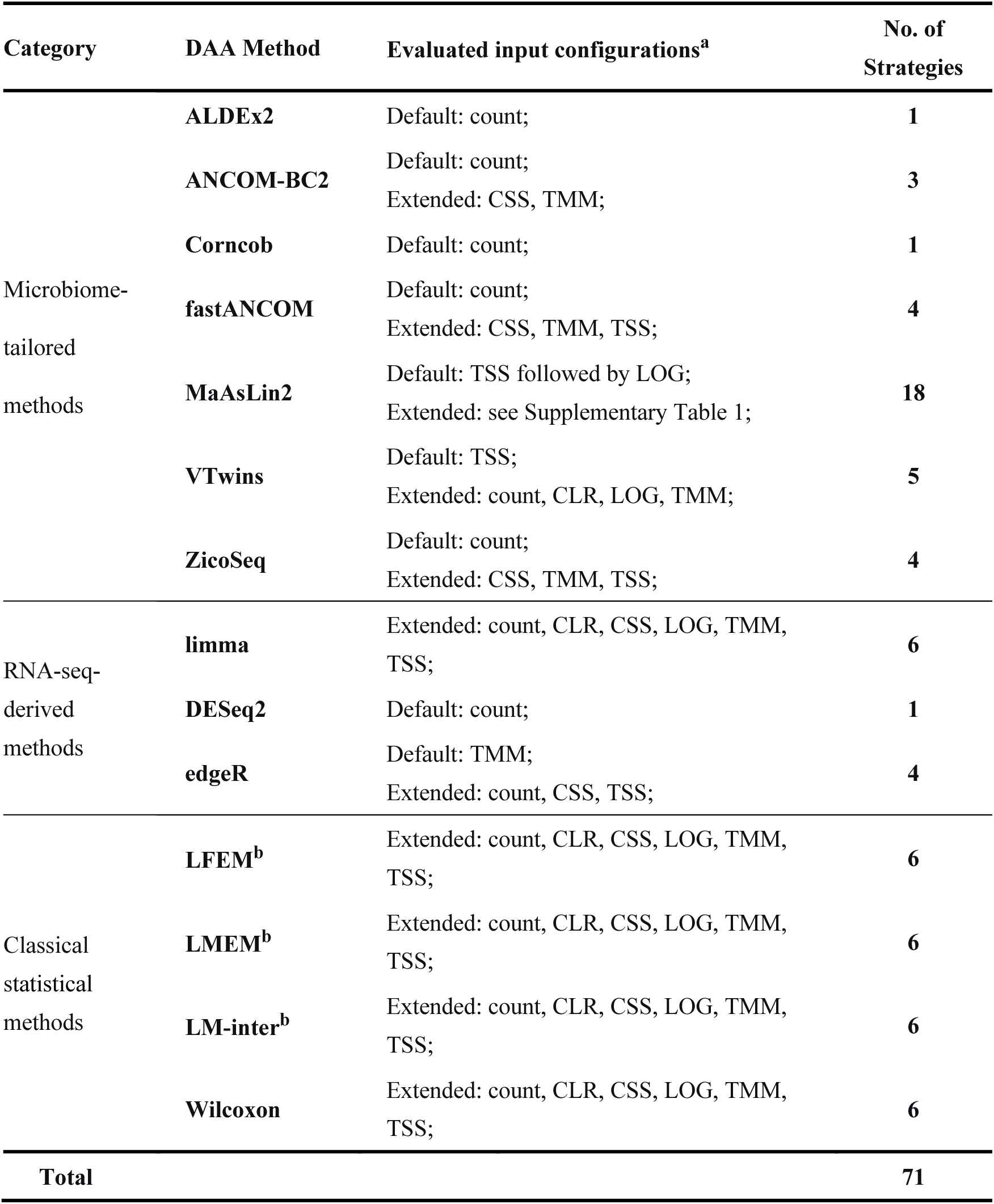
Summary of the 71 benchmarked DAA strategies defined by combinations of data preprocessing procedures and DAA methods.

The single-cohort benchmark followed a staged, multidimensional design (Fig. 1b). All 71 strategies were first evaluated for simulation-based accuracy under known differential abundance signals and predefined confounding structures, together with reference-based recovery of established ecological patterns and disease-associated taxa in oral, vaginal and Crohn’s disease (CD) datasets. Strategies showing persistently weak performance were excluded before resampling-based assessment of within-dataset consistency across 50 disease–control contrasts from 36 gut metagenomic cohorts. Prioritized strategies were subsequently subjected to focused stress testing under systematic perturbations of data characteristics and study-design factors. This staged design generated broad comparative evidence across the 71 strategies and more complete multidimensional performance profiles for the prioritized candidates, supporting interpretation according to analytical objectives rather than a universal ranking (Fig. 1b). We next combined benchmark-prioritized single-cohort strategies with compatible meta-analysis and mega-analysis frameworks, yielding 63 multi-cohort strategies evaluated for simulation-based accuracy under between-cohort heterogeneity and for empirical cross-cohort consistency (Fig. 1c). Finally, we translated the benchmark evidence in xConDA, an interactive web server that links analytical objectives, study designs and data-integration settings to benchmark-prioritized, confounder-adjusted DAA workflows (Fig. 1d).

The strategies are grouped by their underlying DAA method. Complete implementation details for each strategy are provided in the “Benchmark design and construction of single-cohort DAA strategies” section. ^a^ Evaluated strategies were classified as default configurations following developer-recommended or documented workflows and extended configurations representing additional preprocessing–model combinations. ^b^ LFEM, LMEM, and LM-inter denote the linear fixed-effects model, linear mixed-effects model, and linear model with interaction term, respectively.

### 2 Simulation benchmarking of confounder-aware DAA strategies

We first assessed whether explicit confounder adjustment provides measurable benefits under controlled conditions. Using 50 confounded scenarios generated from an IBD-informed simulation framework, we compared adjusted and unadjusted versions of 66 matched DAA strategies in which adjustment status could be changed without altering the underlying statistical model (Supplementary Fig. 1a; Table 1; Supplementary Tables 1 and 2). Covariate adjustment consistently improved error control: all evaluated strategies showed lower false-positive rates (FPR; *P* =1.68 × 10^−12^), and 93.94% showed lower FDR after adjustment (*P* = 1.98 × 10^−11^). This improvement was accompanied by reduced sensitivity (*P* =1.68 × 10^−12^), reflecting a trade-off between suppressing confounding-associated false discoveries and retaining detection power. Threshold-free discrimination was largely preserved, with comparable AUPR (*P* =0.242) and modestly improved AUROC (*P* =5.56 × 10^−3^) after adjustment (Fig. 2a; Supplementary Table 3).

**Fig. 2.**
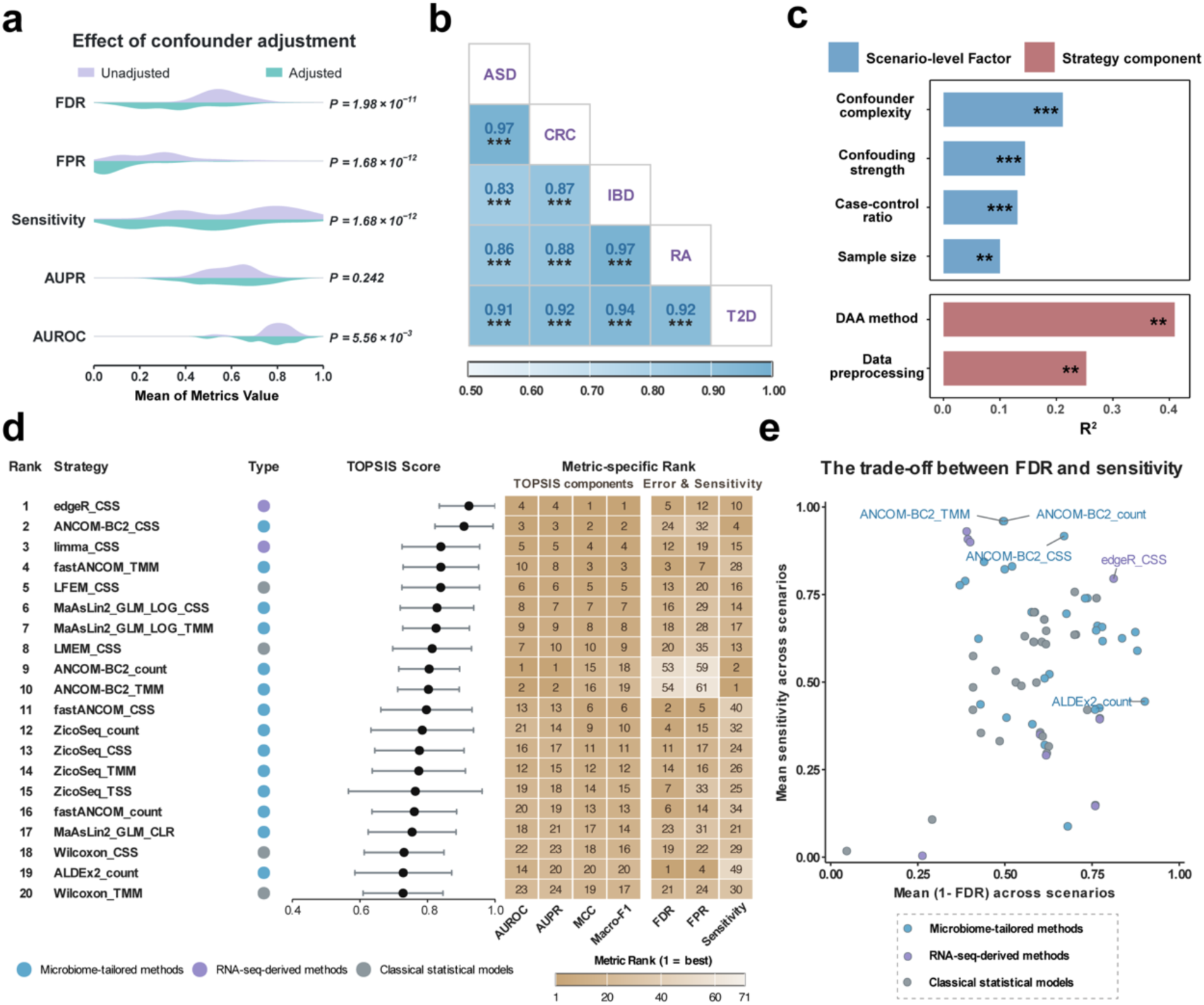
Simulation-based accuracy benchmark of confounder-aware DAA strategies. a, Effect of covariate adjustment on DAA performance. Half-violin plots show the distributions of strategy-level mean FDR, FPR, sensitivity, AUPR and AUROC for 66 matched unadjusted and covariate-adjusted strategies across 50 confounded simulations. Paired differences were assessed by two-sided paired Wilcoxon signed-rank tests. **b**, Consistency of TOPSIS-based strategy rankings across simulations parameterized to autism spectrum disorder (ASD), colorectal cancer (CRC), inflammatory bowel disease (IBD), rheumatoid arthritis (RA) and type 2 diabetes (T2D). Lower-triangle values show pairwise Spearman correlations, with asterisks indicating significance. **c**, Explained variation in performance. PERMANOVA quantified contributions of scenario-level factors and strategy components to variation in integrated TOPSIS scores; bars show the proportion of explained variance (R²). **d**, Composite accuracy ranking and metric-specific profiles of the top 20 strategies. Strategies are ordered by integrated TOPSIS score; black points and horizontal bars show the mean ± s.d. of TOPSIS scores. Colors denote microbiome-tailored, RNA-seq-derived or classical statistical methods. The heatmap shows metric-specific ranks among all 71 strategies. AUROC, AUPR, MCC and Macro-F1 were integrated into the TOPSIS score, whereas FDR, FPR and sensitivity were evaluated separately as operating characteristics; lower values indicate better performance. **e**, Relationship between FDR control and sensitivity. Each point represents one strategy positioned by mean 1 − FDR and mean sensitivity across simulations; upper-right positions indicate a more favorable balance. Significance: no symbol, *P* ≥ 0.1; ·, 0.05–0.1; *, 0.01–0.05; **, 0.001–0.01; ***, *P* < 0.001.

We next systematically benchmarked all 71 end-to-end single-cohort DAA strategies across 250 real-data-informed simulated scenarios, each repeated ten times, yielding 2,500 simulated datasets. Phenotype-associated effects were introduced into multiple microbial features, inducing substantial compositional perturbation and secondary relative-abundance shifts among features without direct phenotype effects. This provided a stringent setting for distinguishing directly perturbed features from secondary compositional shifts [15, 16]. The scenarios spanned five representative microbiome-associated disease contexts—autism spectrum disorder (ASD), CRC, inflammatory bowel disease (IBD), rheumatoid arthritis (RA) and T2D—and incorporated variation in sample size, case–control balance, confounder complexity and confounding strength (Supplementary Fig. 2a, Supplementary Table 2). By preserving key distributional properties of real microbiome datasets while incorporating prespecified differential-abundance signals and confounding structures, these simulations enabled controlled evaluation of strategy performance across diverse conditions. Strategy performance was assessed using complementary metrics, including FPR, FDR, sensitivity, AUROC, AUPR, Macro-F1 and MCC. Replicate-level performance metrics were first averaged within each simulation scenario. AUROC, AUPR, MCC and Macro-F1 were then integrated using the Technique for Order Preference by Similarity to Ideal Solution (TOPSIS) to generate a scenario-specific TOPSIS accuracy score for each strategy. Overall strategy performance was summarized by the mean TOPSIS score across all 250 scenarios, whereas FPR, FDR and sensitivity were examined separately to retain information on error control and detection power (Supplementary Fig. 2b; Supplementary Table 4).

Absolute performance varied across simulation scenarios, demonstrating that DAA accuracy was sensitive to study characteristics and confounding structures (Supplementary Fig. 3). Despite this scenario-level variation, strategy rankings were highly concordant across the five disease-derived simulation templates, with pairwise Spearman correlations ranging from 0.83 to 0.97 (Fig. 2b). Thus, the principal performance patterns were not driven by a single disease-derived dataset. We next examined the factors underlying performance variation. Permutational multivariate analysis of variance (PERMANOVA)-based variance partitioning identified confounder complexity as the largest scenario-level contributor to composite accuracy, followed by confounding strength (Fig. 2c). Among analytical choices, the DAA method accounted for more performance variance (R² = 40.95%, *P* = 0.003) than preprocessing (R² = 25.28%, *P* = 0.007), but preprocessing still substantially affected the performance of individual methods (Supplementary Fig. 3, Fig. 2c). Accordingly, the same DAA method could show markedly different performance when applied to different input representations, and the effect of a preprocessing procedure varied among method families, as illustrated by the contrasting performance of edgeR_CSS and edgeR_TSS (Fig. 2d, Supplementary Fig. 3). These findings support evaluation at the level of complete analytical strategies rather than of preprocessing procedures or DAA methods in isolation.

Mean TOPSIS scores revealed a broad performance gradient among the 71 strategies. edgeR_CSS and ANCOM-BC2_CSS were among the highest-ranked configurations, while other CSS- and TMM-based strategies also showed strong composite performance (Supplementary Fig. 2b, Fig. 2d). However, performance remained multidimensional, and the leading strategies did not dominate across all accuracy metrics. FDR control was particularly scenario-dependent and none of the evaluated strategies achieved an FDR ≤0.05 in all 250 simulation scenarios. ALDEx2_count achieved this threshold most frequently (134/250, 53.6%), but its sensitivity was comparatively low. ANCOM-BC2_TMM and ANCOM-BC2_count showed the contrasting profile, combining higher sensitivity with weaker error control. Across the evaluated strategies, some with relatively low FDR also had comparatively low sensitivity, whereas several highly sensitive strategies showed higher FDR. These method-specific performance profiles illustrate a trade-off between error control and sensitivity (Fig. 2e). These distinct operating profiles argue against reducing strategy evaluation to a single universal ranking and instead motivate selection according to the analytical objective.

We further conducted a targeted sensitivity analysis to assess the robustness of the main performance patterns to signal prevalence. In the sparse-signal setting, 5% of microbial features were prespecified as differentially abundant, compared with 20% in the primary simulations. Using the same TOPSIS framework, we calculated mean strategy scores across the sparse-signal scenarios and compared them with those from the primary simulations. Strategy performance was broadly concordant between the two settings (Spearman’s ρ = 0.766, *P* = 6.5×10^-15^; Supplementary Fig. 4), indicating that the principal performance patterns were largely preserved under lower signal prevalence. ANCOM-BC2_CSS remained among the top-performing strategies, whereas MaAsLin2_CPLM_TSS improved in relative rank under sparse-signal conditions, showing that individual strategy rankings could nevertheless shift with the proportion of differential features (Supplementary Table 4).

Overall, this large-scale simulation show that confounder adjustment, preprocessing and statistical modelling jointly shape DAA performance, while study context can further modify the relative advantages of individual strategies. These findings support benchmarking and selecting complete analytical strategies rather than statistical methods in isolation.

### 3 Reference-based evaluation reveals complementary strategy performance beyond simulation accuracy

Simulation benchmarks enable controlled evaluation against prespecified differential-abundance signals and confounding structures. We therefore complemented these analyses with reference-based evaluations using two complementary frameworks. Reference-pattern recovery was assessed in a paired oral dataset and a vaginal dataset, whereas reference taxa recovery was evaluated in four independent Crohn’s disease (CD) datasets (Fig. 3a). As empirical microbiome datasets lack absolute taxon-level ground truth, these analyses quantified reference-consistent signal recovery rather than absolute accuracy [19].

**Fig. 3.**
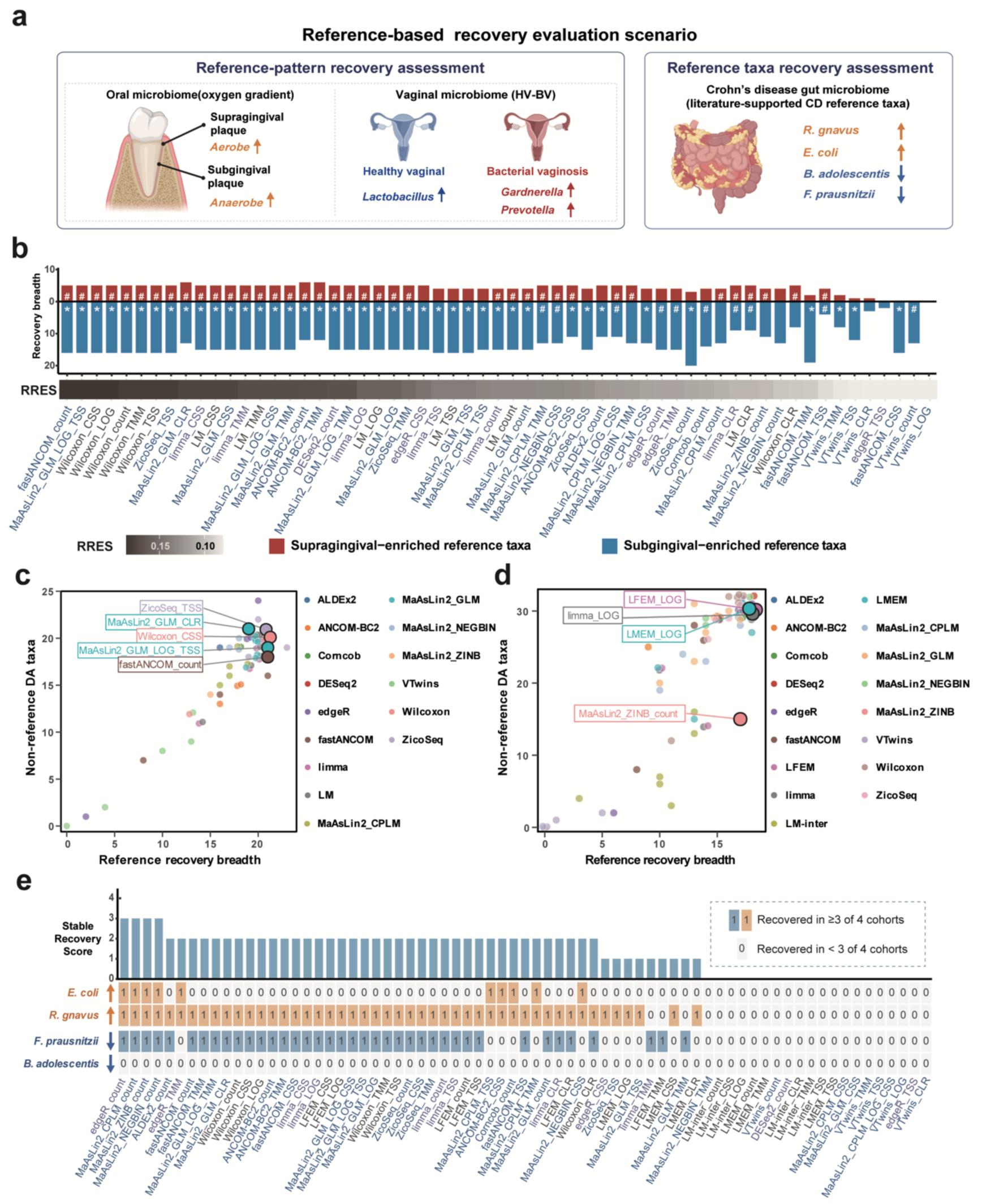
Reference-based recovery evaluation of DAA strategies in empirical microbiome datasets. **a**, Schematic overview of the reference-based evaluation framework. Two complementary settings were performed. Reference-pattern recovery was evaluated in oral and vaginal 16S rRNA datasets using biologically informed microbial patterns. Cross-cohort reference-taxa recovery was assessed using curated CD-associated taxa across four independent CD shotgun metagenomic datasets. **b**, Reference-pattern recovery in the oral microbiome benchmark. Bars show the recovery breadth, defined as the number of reference taxa detected in the expected direction for supragingival-associated aerobes and subgingival-associated anaerobes. Bars are plotted in opposite directions for visualization, with bar magnitude representing recovery breadth. Symbols indicate the significance of reference enrichment among directionally concordant differentially abundant (DA) calls, assessed using one-sided Fisher’s exact tests relative to background recovery: no symbol, *P* > 0.1; #, *P* < 0.1; ∗, *P* < 0.05; ∗∗, *P* < 0.01; ∗∗∗, *P* < 0.001. The lower strip shows the reference recovery enrichment score (RRES), which integrates recovery breadth and reference-enrichment significance; strategies are ordered by decreasing RRES. **c, d**, Relationship between recovery breadth and non-reference DA calls in the oral (**c**) and vaginal (**d**) benchmarks. Each point represents one strategy, positioned by the number of reference taxa recovered in the expected direction and the number of directionally concordant DA taxa outside the corresponding reference set. Points are colored by the DAA strategy family; selected strategies are labelled for reference. **e**, Cross-cohort recovery of curated CD-associated taxa. Bars show the stable recovery score (SRS), defined as the number of the four curated CD reference taxa detected in the expected direction in at least three of the four independent cohorts. The matrix indicates whether each taxon met this stable-recovery criterion for each strategy. Colored cells indicate recovery in the reported direction in at least three cohorts, whereas white cells indicate that the criterion was not met and do not imply absence of recovery in all cohorts. Strategies are ordered by decreasing SRS.

#### Reference-pattern recovery in oral and vaginal microbiome

To evaluate reference-pattern recovery, we considered two complementary aspects of performance: recovery breadth, defined as the number of reference taxa detected in the expected direction, and reference enrichment significance, which assessed whether reference taxa were statistically overrepresented among all directionally concordant differential-abundance calls. Reference enrichment distinguished strategies that selectively recovered the expected pattern from those that achieved high recovery breadth primarily by generating large, nonspecific discovery sets. These two components were integrated into the Reference Recovery Enrichment Score (RRES), which jointly captures the extent of reference recovery and the supporting significant enrichment evidence.

In the oral microbiome, we evaluated recovery of the expected oxygen-associated pattern between paired supragingival and subgingival plaque samples, with aerobic taxa expected to be enriched at supragingival sites and anaerobic taxa at subgingival sites (Fig. 3a). ZicoSeq_count recovered 23 of the 32 predefined reference taxa in the expected direction, the highest recovery breadth among all strategies. fastANCOM_count, MaAsLin2_GLM_LOG_TSS, ZicoSeq_TSS and the Wilcoxon-based family showed comparable taxon-level recovery, each detecting 21 prior taxa. Pattern-level enrichment revealed directional asymmetry. The subgingival anaerobe-enrichment pattern was more readily detected, with 42 strategies showing significant enrichment of anaerobic reference taxa (BH-adjusted *P* < 0.05). For the supragingival aerobe-enrichment pattern, 38 strategies met the more permissive enrichment threshold (BH-adjusted *P* < 0.10) (Supplementary Fig. 5a, Fig. 3b, Supplementary Table 5). Thus, a high overall number of recovered reference taxa did not necessarily imply strong recovery in both reference directions.

After integrating recovery breadth and enrichment strength, fastANCOM_count, ZicoSeq_TSS, the Wilcoxon-based strategies, MaAsLin2_GLM_LOG_TSS and MaAsLin2_GLM_CLR were among the highest-ranked strategies by RRES (Fig. 3b, Supplementary Table 5). Several simulation-leading strategies, including ANCOM-BC2–based strategies, limma_CSS, and LM_CSS, also achieved high RRES values, suggesting partial concordance between simulated accuracy and empirical reference-pattern recovery. In contrast, VTwins-based strategies and fastANCOM_CSS generally showed lower RRES, largely because of limited supragingival-aerobe recovery or weak enrichment evidence in one or both directions (Fig. 3b, Supplementary Table 5). The principal oral-benchmark rankings were broadly preserved across alternative enrichment thresholds, although relaxing the cutoff increased the number of recovered reference taxa (Supplementary Fig. 5b, c). Strategies with broader reference recovery also tended to identify more non-reference taxa, reinforcing the need to interpret recall-like recovery together with enrichment evidence (Fig. 3c). Detectability varied across individual reference taxa: *Fusobacterium*, *Treponema*, *Prevotella*, and *Bacteroides* were readily detected by most strategies, whereas the recovery of *Butyrivibrio* and *Simonsiella* was more strategy-dependent (Supplementary Fig. 6).

We next applied the same framework to vaginal microbiome datasets using 20 curated taxa associated with healthy vaginal (HV) communities or bacterial vaginosis (BV) (Fig. 3a). Reference-pattern recovery varied markedly across strategies, with the number of reference taxa recovered in the expected direction ranging from 0 to 18 (Supplementary Fig. 7a, b, Supplementary Table 6). Recovery also differed between the two phenotype-associated reference sets. Significant enrichment was observed for HV-associated taxa across more strategies, whereas significant enrichment of BV-associated taxa was observed in fewer strategies. Many strategies detected some BV-associated taxa but also identified numerous non-reference taxa, diluting the enrichment signal and reducing statistical significance. MaAsLin2_ZINB_count achieved the highest overall RRES in the vaginal benchmark, combining broad reference recovery with significant enrichment of both the HV- and BV-associated reference sets, while limiting non-reference calls relative to other strategies with similarly broad recovery (Supplementary Fig. 7a, b, Fig. 3d). Several limma-, LFEM- and LMEM-based configurations also achieved favorable RRES values. By contrast, VTwins-based strategies and several TMM-based strategies generally showed narrower reference-pattern recovery, with absent enrichment evidence across the two directions (Supplementary Fig. 7a, b, Supplementary Table 6).

#### Reference taxa recovery assessment in CD microbiome

For disease settings without a predefined ecological enrichment pattern, we evaluated whether externally supported disease-associated taxa could be reproducibly recovered across independent cohorts. We specifically focused on CD, using four independent CD datasets as evaluation cohorts (Fig. 3a). To avoid information leakage, reference taxa were curated exclusively from external CD cohorts that did not overlap with the evaluation datasets. Cross-study synthesis identified four taxa with repeatedly reported, directionally concordant associations with CD (detailed in Methods): *Faecalibacterium prausnitzii* and *Bifidobacterium adolescentis*, which were depleted in CD, and *Escherichia coli* and *Ruminococcus gnavus*, which were enriched in CD. Reproducible recovery was quantified using the Stable Recovery Score (SRS), defined as the number of reference taxa identified by a strategy in the expected direction in at least three of the four cohorts.

No strategy achieved an SRS of four, indicating that, for each strategy, at least one of the four well-supported CD-associated taxa failed to meet the reproducible recovery criterion across cohorts (Fig. 3e, Supplementary Fig. 7c, Supplementary Table 7). Nevertheless, recovery breadth varied markedly among DAA strategies. edgeR_count, MaAsLin2_CPLM_count, MaAsLin2_ZINB_count, and MaAsLin2_NEGBIN_count achieved the highest SRS, each stably recovering three out of the four reference taxa. By contrast, edgeR_TSS, VTwins-based strategies, and LM-inter strategies showed limited recovery and failed to stably recover any CD reference taxa (Fig. 3e, Supplementary Fig. 7c, Supplementary Table 7).

Together, the oral, vaginal and CD benchmarks showed that reference recovery varied substantially across analytical strategies and empirical settings. RRES characterized recovery breadth and reference enrichment within individual datasets, whereas SRS captured directionally consistent recovery across independent cohorts. These analyses revealed strategy-specific strengths that were not captured by simulation-based accuracy alone, supporting reference-based recovery as a complementary benchmark dimension.

### 4 Within-dataset consistency and perturbation stability reveal distinct robustness profiles

To complement simulation-based accuracy and empirical reference recovery, we next evaluated two dimensions of strategy robustness: within-dataset consistency under resampling of empirical cohorts and performance stability under systematic perturbation of simulated data characteristics. To focus these analyses on plausible candidate workflows, we removed 11 strategies that were consistently low performing across earlier benchmarks. Strategies were excluded if they ranked in the bottom 25% in at least two reference-based benchmarks and in the bottom 50% in both the overall simulation-based accuracy benchmark and the sparse-signal sensitivity analysis, leaving 60 strategies for within-dataset consistency assessment. A subsequent consistency-based filter further prioritized 12 strategies for perturbation stability analysis (Supplementary Fig. 8).

#### Within-dataset consistency of differential feature sets

Within-dataset consistency was evaluated across 50 disease-control datasets derived from 36 human gut metagenomic cohorts by comparing differentially abundant (DA) feature sets obtained from different sample subsets of the same contrast. The Jaccard index (JI) quantified consistency between subset-derived DA feature sets, while the number of detected DA features captured detection scale and helped distinguish consistent recovery from overly sparse outputs.

Consistency varied substantially across strategies and empirical datasets (Fig. 4a, Supplementary Table 8). MaAsLin2 negative binomial models (NEGBIN) showed the strongest overall within-dataset consistency, with MaAsLin2_NEGBIN_TMM and MaAsLin2_NEGBIN_count achieving the highest mean JI values while maintaining relatively high mean number of detected DA features. Conversely, strategies, such as limma_CSS, ZicoSeq_TSS, and fastANCOM_count, frequently yielded low or zero JI in many individual datasets, indicating limited within-dataset consistency of DA feature recovery under resampling.

**Fig. 4.**
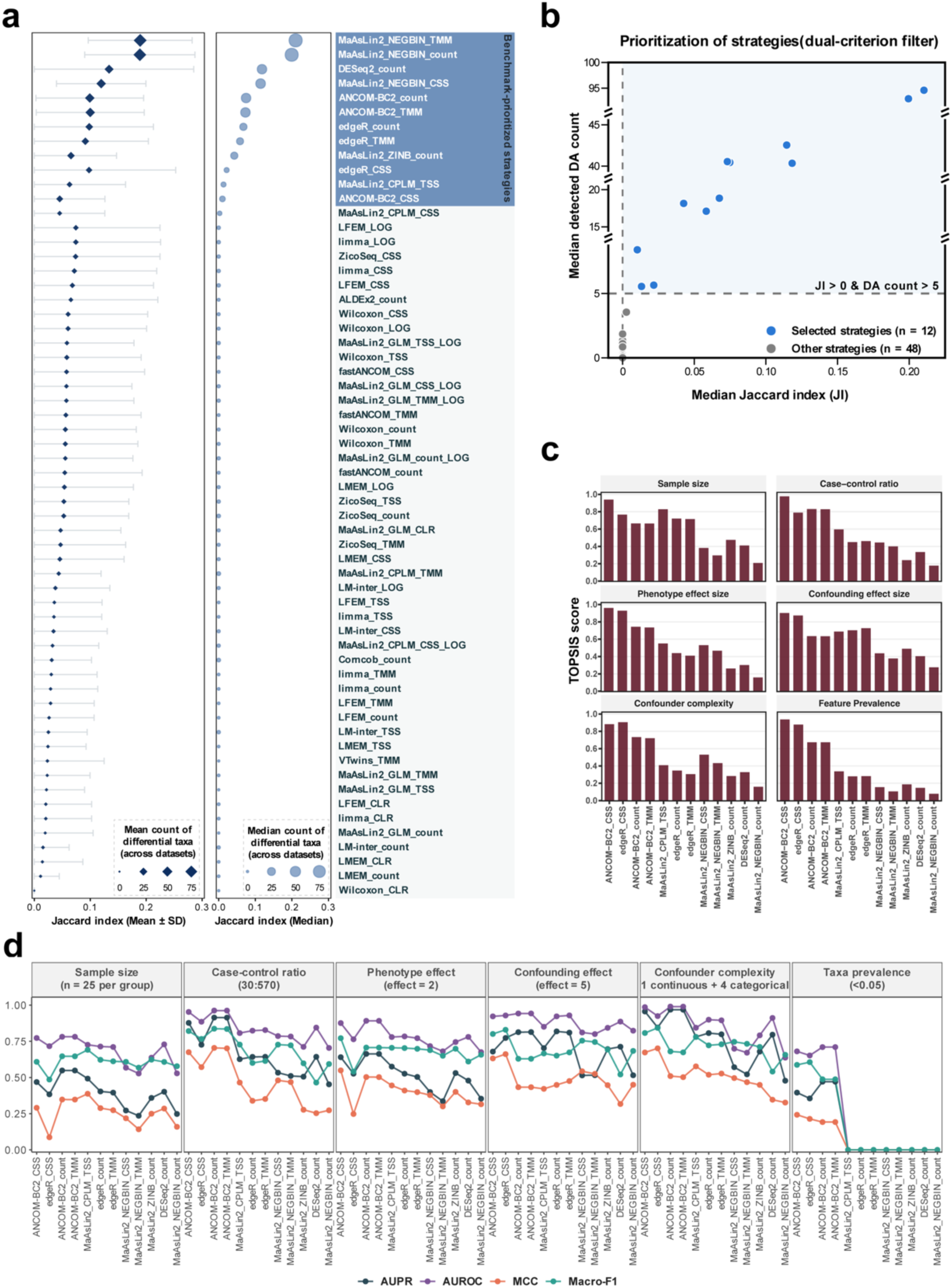
Robustness evaluation and prioritization of confounder-aware DAA strategies. **a**, Within-dataset consistency across 50 disease–control contrasts from 36 human gut metagenomic cohorts. For each strategy, Jaccard index (JI) was calculated between sets of DA taxa identified from paired resampled subsets within each dataset. Dark-blue diamonds and horizontal bars show the mean ± s.d. of JI across evaluated datasets, with diamond size representing the mean number of detected DA taxa. Light-blue circles show the median JI, with circle size representing the median number of detected DA taxa. The blue-shaded label block indicates the 12 benchmark-prioritized strategies. **b**, Strategy prioritization by dual-criterion filtering. Each point represents one of the 60 strategies evaluated for within-dataset consistency. Strategies satisfying both a median JI > 0 and a median detected DA count > 5 across subset comparisons were prioritized. Dashed lines indicate the selection thresholds, and the shaded region denotes the criterion space. Prioritized strategies are shown in blue (*n* = 12) and the remaining strategies in grey (*n* = 48). Median detected DA counts are displayed on a broken y-axis, with zero values placed at the lower display boundary. **c,** Perturbation robustness of the 12 benchmark-prioritized strategies across six perturbation factors. For each factor, AUROC, AUPR, MCC, and Macro-F1 were evaluated across perturbation levels and summarized by the AUMTC. The four metric-specific AUMTC values were integrated by TOPSIS into a composite score. Higher scores indicate stronger overall performance across perturbation levels. **d,** Performance of prioritized strategies under predefined challenging perturbation settings. For each perturbation factor, AUROC, AUPR, MCC and Macro-F1 were evaluated at the challenging level indicated in the facet header. The evaluated factors were sample size, case–control ratio, phenotype effect size, confounding effect size, confounder complexity and feature prevalence. Full parameter settings are provided in Supplementary Table 12.

To prioritize strategies showing consistent feature recovery without overly sparse discovery sets, we applied a dual-criterion filter based on both reproducibility and detection scale, requiring a median detected DA count greater than 5 and a median JI greater than 0 across subset comparisons (Fig. 4b, Supplementary Fig. 8). This retained 12 benchmark-prioritized single-cohort strategies, including MaAsLin2_NEGBIN_TMM, MaAsLin2_NEGBIN_count, MaAsLin2_NEGBIN_CSS, ANCOM-BC2_TMM, ANCOM-BC2_count, DESeq2_count, edgeR_count, MaAsLin2_ZINB_count, edgeR_TMM, ANCOM-BC2_CSS, edgeR_CSS, and MaAsLin2_CPLM_TSS (Fig. 4a).

#### Performance stability across systematic perturbations

We next evaluated the 12 benchmark-prioritized strategies under systematic perturbations of six key data and study-design factors: sample size, case–control ratio, phenotype effect size, confounding feature effect size, confounder complexity and feature prevalence (Supplementary Table 9, detailed in Methods). For each perturbation factor, AUROC, AUPR, MCC and Macro-F1 were tracked across factor levels and summarized using the area under the metric–trend curve (AUMTC) (Fig. 4c, Supplementary Fig. 9a; detailed in Methods). Metric-specific AUMTC values were then integrated using TOPSIS to summarize sustained performance across perturbation levels and obtain an overall strategy ranking.

ANCOM-BC2_CSS and edgeR_CSS showed the strongest overall perturbation-robustness profiles, ranking consistently among the top-performing strategies across changes in sample size, case–control balance, phenotype effect size, confounding effect size, confounder complexity and feature prevalence (Fig. 4c, Supplementary Fig. 9 and Supplementary Table 10). Other strategies showed more context-specific strengths, for example, MaAsLin2_CPLM_TSS performed competitively under sample-size perturbation, whereas ANCOM-BC2_TMM and ANCOM-BC2_count performed well across several design-related perturbations, including case–control ratio, phenotype effect size, confounder complexity, and feature prevalence, but ranked just below the top-performing strategies in overall robustness.

Metric-specific trajectories provided further insight into these differences. The broad robustness of ANCOM-BC2_CSS and edgeR_CSS was driven primarily by stable error control rather than uniformly high sensitivity, a property particularly relevant for conservative inference in confounded microbiome studies (Supplementary Fig. 9a, Supplementary Table 11a). Feature prevalence perturbation was a notable exception, under which both strategies maintained high sensitivity while preserving strong FDR control (Supplementary Fig. 9a). ANCOM-BC2_TMM and ANCOM-BC2_count maintained strong AUROC/AUPR and high sensitivity across several perturbations but showed less stable MCC and Macro-F1, resulting in lower integrated robustness rankings, with their lower integrated robustness largely reflecting less stable FDR control rather than reduced detection sensitivity (Supplementary Fig. 9a, Supplementary Tables 10 and 11). FPR broadly paralleled FDR and provided limited additional discrimination among strategies; it is therefore not discussed further.

Worst-case analyses under the most challenging perturbation levels were consistent with the AUMTC-based results (Fig. 4d, Supplementary Table 12). ANCOM-BC2_CSS maintained strong performance across all challenging settings examined. edgeR_CSS also retained favorable performance under extremely imbalanced case– control ratios, strong confounder effects, complex confounding structures, and low feature prevalence. Beyond the globally stable strategies, several strategies also showed scenario-specific strengths, for example, MaAsLin2_CPLM_TSS performed well under small-sample settings, while ANCOM-BC2_TMM/count excelled in small sample size, extreme case–control imbalance, and weak phenotype effect sizes (Fig. 4d, Supplementary Table 12).

These analyses revealed that within-dataset consistency and perturbation robustness capture complementary properties of DAA strategies. MaAsLin2_NEGBIN_TMM and MaAsLin2_NEGBIN_count favored reproducible DA feature recovery under resampling, whereas ANCOM-BC2_CSS and edgeR_CSS showed broader robustness to systematic perturbations. These distinct profiles support objective-guided strategy prioritization.

### 5 Multidimensional benchmarking enables objective-guided single-cohort strategy selection

We integrated evidence across four dimensions of single-cohort benchmark— simulation-based accuracy, empirical reference recovery, within-dataset consistency and perturbation stability—to characterize the multidimensional performance profiles of the 12 benchmark-prioritized strategies (Fig. 5a). Rather than defining a universally optimal workflow, this integration framework summarizes strategy-specific strengths and trade-offs across distinct analytical objectives. Performance across the four benchmark dimensions was summarized into an overall integrated score to facilitate comparison and visualization of multidimensional performance profiles, while dimension-specific metrics were retained to reveal objective-dependent differences among strategies.

**Fig. 5.**
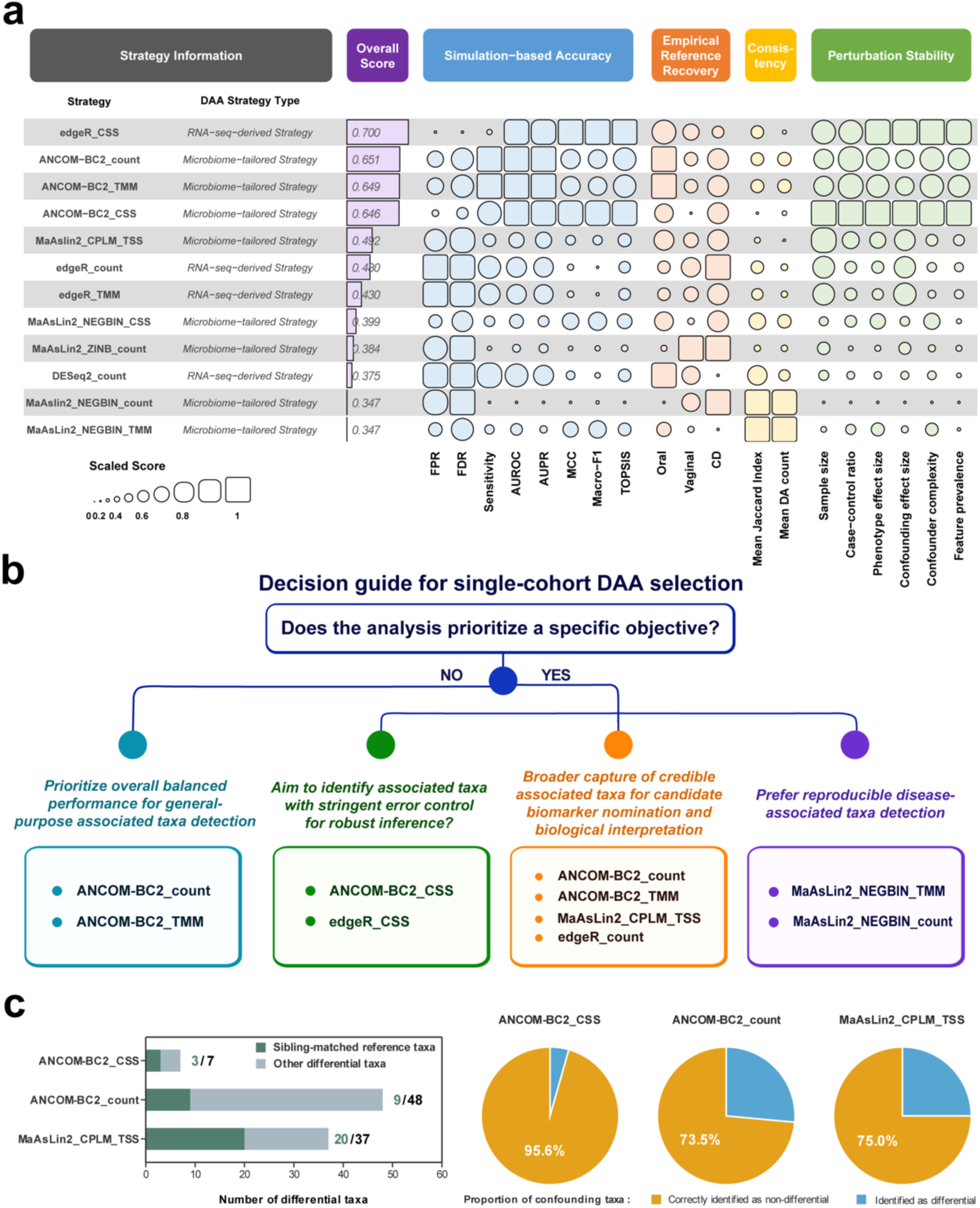
Integrated single-cohort benchmarking supports objective-guided DAA strategy selection. a, Multidimensional performance profiles of the 12 benchmark-prioritized single-cohort DAA strategies, ordered by an overall integrated score that summarizes performance across four benchmark dimensions, as described in Methods. Columns summarize DAA strategy category and performance across simulation-based accuracy, reference-based recovery, within-dataset consistency and perturbation stability. Simulation-based accuracy includes FPR, FDR, sensitivity, AUROC, AUPR, MCC, Macro-F1, and the integrated TOPSIS score. Empirical reference-based recovery summarizes the oral oxygen-associated, vaginal HV–BV and Crohn’s disease reference-taxa benchmarks. Within-dataset consistency is summarized by mean Jaccard index and mean detected DA taxa, whereas perturbation stability performance is summarized by TOPSIS-integrated AUMTC scores across six perturbation factors. Displayed metrics were scaled to 0–1, with larger symbol size indicating better relative performance. **b,** Objective-guided decision framework for single-cohort DAA strategy selection. For general-purpose analyses without a specific objective, strategies with balanced overall performance are prioritized. For analyses with a specific objective, strategies are prioritized according to the primary analytical goal: stringent error control for high-confidence inference; broader signal recovery for candidate discovery and biological interpretation; reproducible detection of associated taxa. **c,** Practical application of benchmark-prioritized strategies in CD case study informed by a sibling-matched design. Stacked bar plots show the total number of DA taxa detected in confounder-adjusted comparisons of active CD versus unrelated controls for ANCOM-BC2_CSS, ANCOM-BC2_count, and MaAsLin2_CPLM_TSS. Dark segments indicate sibling-matched reference taxa and light segments indicate other differential taxa. Values beside the bars denote sibling-matched reference taxa / total detected DA taxa for each strategy. Pie charts show the proportions of predefined putative familial or environment-associated taxa classified as differential or non-differential by each strategy. Orange denotes taxa classified as non-differential, and blue denotes taxa identified as differential.

Using these multidimensional profiles, we developed an objective-guided decision framework for single-cohort DAA strategy selection (Fig. 5b). For general-purpose analyses without a predefined analytical objective, ANCOM-BC2_count and ANCOM-BC2_TMM showed comparatively balanced performance across benchmark dimensions and provided broadly applicable options. When stringent error control was prioritized for high-confidence inference, such as validation-oriented analyses, mechanistic follow-up or prioritization of high-confidence taxa, edgeR_CSS and ANCOM-BC2_CSS were recommended because they combined strong simulation-based accuracy and favorable error control with broad perturbation stability (Fig. 5a, b). For analyses emphasizing broader recovery of credible associated taxa, such as candidate biomarkers or biological interpretation, ANCOM-BC2_count, ANCOM-BC2_TMM, MaAsLin2_CPLM_TSS, and edgeR_count provided discovery-oriented options with favorable empirical reference-recovery profiles. When reproducible feature detection under sample resampling is the primary goal, MaAsLin2_NEGBIN_TMM and MaAsLin2_NEGBIN_count showed the strongest within-dataset consistency (Fig. 5a, b). Sample-size-stratified results also suggested that ANCOM-BC2_count and ANCOM-BC2_TMM are favored for small-sample studies because their stronger discriminative performance and relatively high sensitivity supports signal detection under limited statistical power. For larger studies where sensitivity is generally high, edgeR_CSS is favored for its comparatively strong FDR control (Supplementary Tables 11 and 12).

To illustrate benchmark-guided strategy selection in practice, we applied representative strategies from different decision branches to an active CD case study informed by a sibling-matched design (Fig. 5c, Supplementary Table 13). Unaffected healthy siblings provide a family-matched comparison that partially controls for shared genetic and environmental background. Using our previous sibling-matched CD study [8], we evaluated strategy performance using sibling-matched reference taxa representing putative disease-associated signals and family-design-derived confounding-sensitive taxa representing potential familial or environmental influences. Using an adjusted *P*-value threshold of 0.1, ANCOM-BC2_count recovered nine sibling-matched reference taxa, while 73.5% of the predefined confounding-sensitive taxa remained non-differential. MaAsLin2_CPLM_TSS showed broader signal recovery, identifying 20 sibling-matched reference taxa, while 75.0% of the confounding-sensitive taxa remained non-differential. In contrast, ANCOM-BC2_CSS prioritized conservative inference, recovering fewer sibling-matched reference taxa (three taxa), while 95.6% of the predefined confounding-sensitive taxa remained non-differential. This conservative behavior was maintained at a relaxed threshold of adjusted P of 0.2, at which ANCOM-BC2_CSS still excluded 92.6% of the predefined confounding-sensitive taxa. These results illustrate different operating characteristics among benchmark-prioritized strategies, with broader recovery favoring exploratory discovery and stricter control limiting potentially background-associated signals. More broadly, the integrated benchmark maps multidimensional performance onto explicit analytical objectives, supporting objective-guided selection of complete DAA strategies rather than a universal method ranking.

### 6 Multi-cohort integration reshapes DAA performance and strategy priorities

We next extended the benchmark to multi-cohort DAA, where between-cohort heterogeneity, batch effects and incomplete covariate harmonization introduce additional analytical challenges. Each multi-cohort strategy combined a benchmark-prioritized single-cohort preprocessing–model configuration with a cross-cohort integration mode. Meta-analysis applied DAA separately within each cohort and combined cohort-level statistics [30], whereas mega-analysis pooled individual-level data and modeled cohort as an adjustment factor [28]. Combining the 12 benchmark-prioritized single-cohort strategies with compatible integration frameworks yielded 63 multi-cohort DAA strategies (Fig. 6a; detailed in Methods, Supplementary Table 14). In multi-cohort simulations with pronounced cohort-level bias and limited harmonizable covariates (detailed in Methods), mega-analysis strategies based on ANCOM-BC2_CSS and edgeR_CSS achieved the highest overall accuracy (Fig. 6b, c, Supplementary Table 15). edgeR_CSS_Mega favored sensitivity, whereas ANCOM-BC2_CSS_Mega achieved stronger error control while maintaining competitive sensitivity. Integration mode also substantially influenced performance. For the same underlying single-cohort DAA strategy, effect-size-based meta-analysis generally outperformed P-value-based aggregation across most accuracy metrics when comparable effect estimates and standard errors were available. Several effect-size-based meta-analysis strategies, including those derived from ANCOM-BC2_CSS and MaAsLin2_CPLM, performed favorably across multiple accuracy dimensions (Supplementary Fig. 10, Supplementary Table 15). By contrast, MaAsLin2_NEGBIN-based strategies ranked lower in the simulation benchmark, particularly under mega-analysis or P-value-based meta-analysis implementations (Supplementary Fig. 10, Supplementary Table 15). Thus, multi-cohort DAA performance depends both on the underlying single-cohort DAA strategy and the mode of multi-cohort integration.

**Fig. 6.**
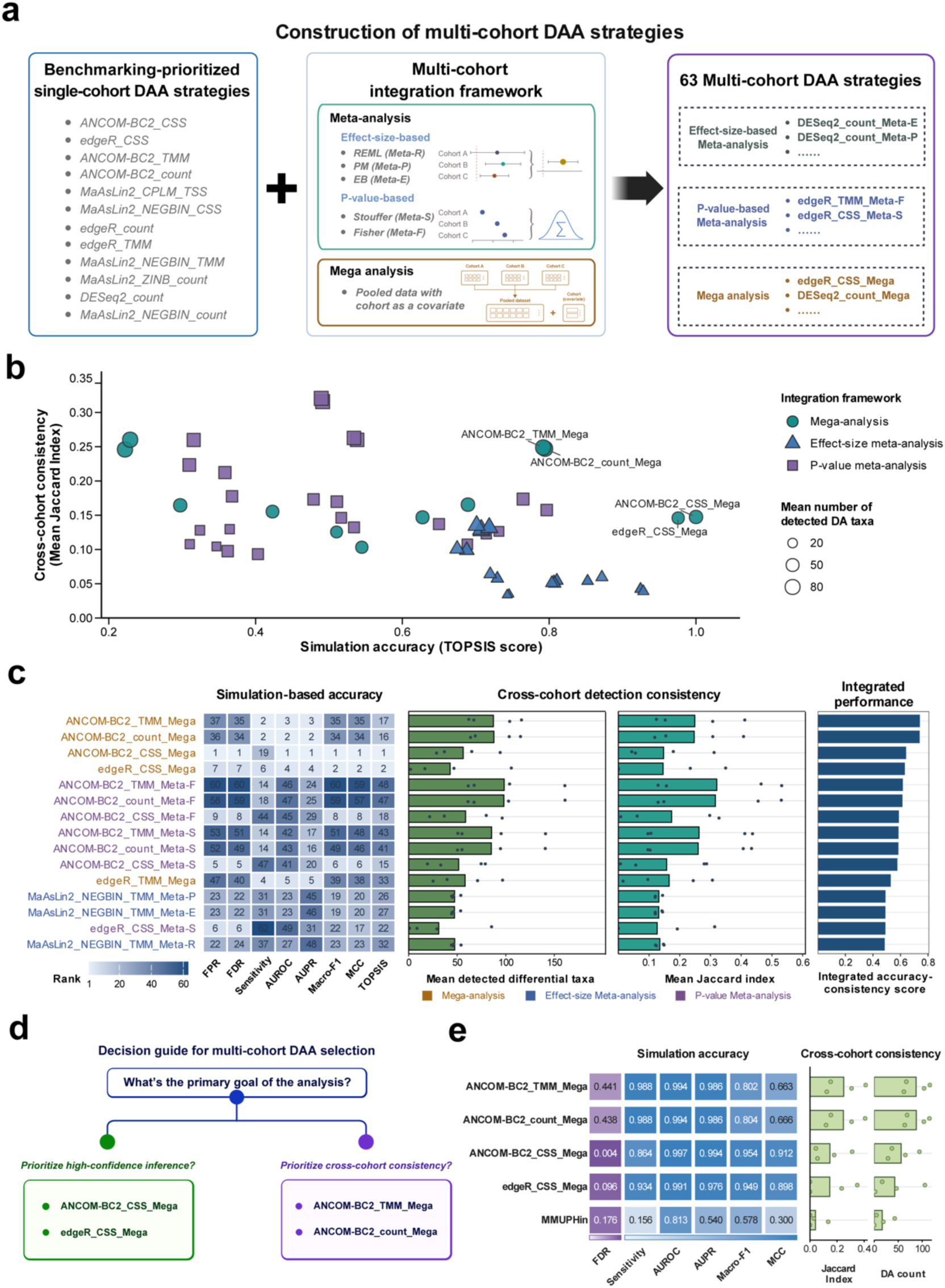
Benchmarking of multi-cohort DAA strategies. **a**, Construction of multi-cohort DAA strategies. The 12 benchmark-prioritized single-cohort strategies were extended using meta-analysis or mega-analysis frameworks. Meta-analysis was further divided into P-value-based and effect-size-based approaches, whereas mega-analysis pooled cohorts with cohort included as a covariate, yielding 63 multi-cohort strategies. **b**, Relationship between simulation-based accuracy and empirical cross-cohort consistency across the 63 strategies. Each point represents one strategy, positioned by its simulation TOPSIS score and mean cross-cohort Jaccard index (JI). Point shape and color denote the integration framework, and point size represents the mean number of detected DA taxa across the empirical disease settings. Selected strategies are labelled. **c**, Integrated performance profiles of the top 15 multi-cohort DAA strategies. The heatmap shows metric-specific ranks among all 63 strategies for simulation-based performance, with lower ranks indicating better performance. The middle panels summarize empirical cross-cohort detection scale and consistency by mean DA count and mean JI, with points showing disease-specific values. The rightmost bars show the integrated accuracy–consistency score, combining the simulation TOPSIS score and mean JI via TOPSIS. Strategies are ordered from highest to lowest integrated score. Strategy-label colors denote the integration framework: gold, mega-analysis; blue, effect-size meta-analysis; and purple, P-value meta-analysis. Complete results for all 63 strategies are shown in Supplementary Fig. 10. **d**, Objective-guided selection of benchmark-prioritized mega-analysis strategies. ANCOM-BC2_CSS_Mega and edgeR_CSS_Mega are favored for high-confidence inference, whereas ANCOM-BC2_TMM_Mega and ANCOM-BC2_count_Mega are favored for cross-cohort consistency. **e**, Comparison of four benchmark-prioritized multi-cohort DAA strategies with MMUPHin. Left, simulation accuracy for FDR, sensitivity, AUROC, AUPR, Macro-F1 and MCC. Cell labels show the original values; darker purple indicates lower FDR, whereas darker blue indicates higher values for the other metrics. Right, real-data consistency based on the mean direction-concordant JI and mean DA count. Light-green bars show means across four disease datasets, and green circles indicate disease-specific values.

We next assessed cross-cohort consistency using multi-cohort metagenomic datasets from curatedMetagenomicData [31], covering adenoma, CRC, IBD, and T2D. For all 63 multi-cohort strategies, detection consistency was quantified from the overlap of significant DA taxa across different cohort groupings, with detection yield assessed in parallel. Several P-value-based meta-analysis strategies, particularly ANCOM-BC2_TMM_Meta-F, ANCOM-BC2_count_Meta-F, ANCOM-BC2_TMM_Meta-S, and ANCOM-BC2_count_Meta-S, together with several MaAsLin2_NEGBIN-based strategies, achieved high cross-cohort consistency and relatively broad DA detection, despite ranking lower in the simulation benchmark (Fig. 6b, c, Supplementary Fig. 10, Supplementary Tables 15 and 16). Conversely, ANCOM-BC2_CSS_Mega and edgeR_CSS_Mega, which ranked among the strongest strategies in the simulation benchmark, showed only moderate detection consistency (Fig. 6b, c, Supplementary Tables 15 and 16).

As a sensitivity analysis, we incorporated effect direction for the 39 strategies providing signed integrated effect estimates. Requiring concordant effect directions across cohort groupings moderately reduced consistency scores but largely preserved the relative strategy rankings (Spearman’s ρ = 0.966, *P* < 2.2 × 10^−16^; Supplementary Fig. 11a, Supplementary Table 17). ANCOM-BC2_count_Mega, ANCOM-BC2_TMM_Mega, MaAsLin2_NEGBIN_count_Mega, and MaAsLin2_NEGBIN_TMM_Mega ranked among the highest for both detection consistency and direction-concordant detection consistency (Supplementary Fig. 11b, Supplementary Table 17). These findings indicated that the principal consistency patterns were robust to requiring concordant association directions.

Joint consideration of simulation accuracy and cross-cohort consistency prioritized four mega-analysis strategies with complementary performance profiles (Fig. 6c). ANCOM-BC2_CSS_Mega and edgeR_CSS_Mega, both built on single-cohort extended input configurations, achieved strong accuracy and error control together with moderate cross-cohort consistency, supporting their use for high-confidence multi-cohort inference. ANCOM-BC2_TMM_Mega and ANCOM-BC2_count_Mega showed higher cross-cohort consistency while maintaining competitive simulation performance, supporting reproducibility-oriented applications (Fig. 6d). Integrating simulation accuracy with direction-concordant consistency among the 39 eligible strategies identified the same four strategies and preserved this objective-specific prioritization (Supplementary Fig. 11). Overall, mega-analysis provided the strongest balance of benchmark dimensions when individual-level data could be jointly modeled, whereas meta-analysis remained a practical alternative when only cohort-level results could be integrated.

We further compared the four prioritized multi-cohort strategies with MMUPHin [32], a widely used tool for multi-cohort microbiome analysis. Under our benchmark settings, the prioritized strategies generally showed stronger integrated simulation performance and higher empirical cross-cohort consistency than MMUPHin (Fig. 6e, Supplementary Tables 15 and 16). Although MMUPHin showed stronger FDR control than ANCOM-BC2_TMM_Mega and ANCOM-BC2_count_Mega, its markedly lower sensitivity led to weaker overall performance across AUROC, AUPR, MCC, and Macro-F1. In the real-data consistency evaluation, MMUPHin also detected fewer differentially abundant species and showed lower cross-cohort reproducibility. These comparisons further highlighted the trade-offs among multi-cohort strategies in error control, detection yield and cross-cohort consistency.

Finally, we examined the four prioritized mega-analysis strategies and MMUPHin in a multi-cohort CRC application. All five strategies recovered a shared core of established CRC-enriched species, including *Fusobacterium nucleatum*, *Parvimonas micra*, *Peptostreptococcus stomatis*, *Solobacterium moorei*, and *Gemella morbillorum* (Supplementary Fig. 12 and Supplementary Table 18). Beyond this shared core, detection breadth differed substantially among strategies. ANCOM-BC2_TMM_Mega and ANCOM-BC2_count_Mega identified broader DA species sets, whereas edgeR_CSS_Mega, ANCOM-BC2_CSS_Mega and MMUPHin were more conservative. Several additional species showed strategy-dependent detection patterns without contradictory effect directions. *Streptococcus thermophilus* was detected by all prioritized mega-analysis strategies but not by MMUPHin, whereas *Dialister pneumosintes*, *Bacteroides fragilis* and *Clostridium symbiosum* were detected by most strategies but not by the more conservative ANCOM-BC2_CSS_Mega workflow. In contrast, *Akkermansia muciniphila* was detected only by MMUPHin and the two broader-detection ANCOM-BC2 mega-analysis strategies. *Alistipes onderdonkii* was not detected by MMUPHin or edgeR_CSS_Mega. Thus, the prioritized strategies and MMUPHin consistently recovered a common CRC-associated signal while differing in the breadth of additional taxa detected.

### 7 xConDA implements benchmark-guided, objective-aware DAA strategy selection

To translate the benchmark into an actionable analysis resource, we developed xConDA (Confounder-aware Differential Abundance Analysis; https://www.biosino.org/xconda/), an interactive web server that translates systematic benchmark results into executable, benchmark-prioritized confounder-aware DAA workflows for single- and multi-cohort microbiome studies. xConDA links strategy recommendation to the user’s analytical objective, including high-confidence inference, biological signal recovery, reproducible feature detection and multi-cohort integration, and displays the corresponding multidimensional benchmark evidence to support strategy prioritization.

Users can upload a microbial abundance table and sample metadata, define the phenotype and covariates, and execute standardized preprocessing, covariate-adjusted DAA and, when applicable, multi-cohort integration. xConDA returns harmonized result tables, visualizations and analysis reports, facilitating standardized comparison and interpretation across heterogeneous DAA strategies.

By integrating benchmark evidence, objective-guided strategy selection and standardized execution, xConDA provides an executable decision-support resource for confounder-aware microbiome DAA in single- and multi-cohort microbiome studies.

## Discussion

Reliable differential abundance analysis remains challenging because microbiome associations are sensitive to confounding and to interacting choices in data preprocessing, statistical modelling and cross-cohort integration. We therefore benchmarked confounder-aware DAA at the level of complete analytical strategies, integrating simulation-based accuracy, empirical reference recovery and robustness across single- and multi-cohort settings. This strategy-level, multidimensional framework extends conventional method-centric comparisons by evaluating analytical choices that are applied together in practice. Covariate adjustment generally strengthened error control at the cost of reduced sensitivity, while different strategies showed distinct strengths across benchmark dimensions. Rather than identifying a universally optimal workflow, our benchmark revealed objective-dependent operating profiles that support strategy selection according to the analytical goal.

A central finding was that covariate adjustment substantially improved false-positive and false-discovery control but reduced sensitivity. This reinforces growing concerns that host, clinical and technical factors can shape apparent microbiome–disease associations [3, 33, 34]. Rigorous covariate control can alter previously reported disease-associated microbial signals [3], and our results show that adjustment changes the balance between error control and signal detection rather than uniformly improving performance. These findings extend recent simulation-based evaluation of confounder adjustment [16] by showing that substantial variation remains after adjustment and depends on the broader analytical strategy. Both the DAA method and preprocessing contributed strongly to the strategy performance, with individual method performance varying substantially across preprocessing choices, consistent with evidence that normalization and transformation can alter microbiome inference [18, 22, 35]. Accordingly, confounder adjustment should be treated as one design component within a complete analytical workflow rather than as an independent solution to confounding. Relevant confounders may also remain unmeasured or misspecified, and statistical adjustment does not establish causality. Reliable DAA therefore requires evaluation of the preprocessing–model–adjustment combination rather than the statistical method alone [15, 16, 18, 19].

Our multidimensional evaluation further shows why simulation-based accuracy alone is insufficient to characterize practical DAA performance [15, 21]. Simulations provide prespecified signals and confounding structures and are therefore essential for evaluating statistical accuracy and error control, but even real-data-informed simulations cannot reproduce the full complexity of empirical microbiome data [36, 37]. Conversely, empirical datasets lack absolute taxon-level ground truth. We therefore complemented simulation with empirical reference recovery to assess whether analytical strategies recover biologically plausible signals, while within-dataset consistency assessed whether similar features were recovered despite within-cohort sample heterogeneity [19, 20]. Perturbation robustness further assessed whether performance was sustained as study characteristics changed. These additional evaluation dimensions were reflected in the strategy rankings: methods favored in previous simulation-based benchmarks, including ZicoSeq [15], limma, fastANCOM and other linear-model-based approaches [16], were not consistently prioritized when empirical reference recovery and reproducibility were considered. The fact that these dimensions did not consistently favor the same strategies indicates that they capture complementary aspects of DAA performance. Their joint consideration provides a more informative basis for objective-guided strategy selection than relying on any single performance dimension alone.

This multidimensionality has a direct practical implication: the preferred strategy depends on the primary analytical objective. ANCOM-BC2_CSS and edgeR_CSS were favored for stringent error control and perturbation-robust, high-confidence inference, whereas MaAsLin2_NEGBIN configurations showed stronger within-dataset consistency. ANCOM-BC2_count and ANCOM-BC2_TMM provided comparatively balanced profiles when no single objective dominated. Notably, ANCOM-BC2_CSS, edgeR_CSS and ANCOM-BC2_TMM were extended input configurations, highlighting the importance of systematically exploring preprocessing–DAA method combinations rather than limiting evaluation to canonical configurations [18, 22, 35]. These differences highlight distinct performance strengths across strategies rather than supporting a single composite ranking. Conservative workflows may be preferable when false discoveries carry substantial downstream costs, whereas broader signal recovery may be appropriate for exploratory analyses followed by independent validation. The sibling-matched CD analysis further illustrates this principle: unaffected siblings partly account for shared familial and environmental exposures [8], and ANCOM-BC2_CSS and MaAsLin2_CPLM_TSS showed distinct trade-offs between restricting potentially confounding-sensitive associations and recovering broader disease-associated signals.

Multi-cohort studies introduce an additional analytical decision because performance depends not only on the underlying DAA strategy but also on how evidence is integrated across cohorts. Between-cohort heterogeneity is widespread in microbiome studies [23–26], and combining cohorts cannot be regarded simply as increasing sample size. In the settings evaluated here, mega-analysis provided the strongest overall balance when harmonizable individual-level data were available. This advantage is conditional on access to compatible individual-level data and should not be interpreted as universal superiority [38, 39]. When individual-level pooling is not feasible, meta-analysis remains an important alternative [27]. Effect-size integration was generally favored when comparable effect estimates and standard errors were available, whereas P-value aggregation remained practical when only limited summary information could be harmonized. Objective dependence also persisted after integration: ANCOM-BC2_CSS_Mega and edgeR_CSS_Mega favored simulation-based accuracy, whereas ANCOM-BC2_TMM_Mega and ANCOM-BC2_count_Mega favored cross-cohort consistency. Comparison with MMUPHin [32] further illustrated that multi-cohort approaches differed in their performance strengths, with no universally optimal strategy. These results provide practical guidance for selecting multi-cohort analysis strategies according to data availability and analytical priorities.

Benchmark evidence is most useful when it can be translated into reproducible analytical decisions. We therefore developed xConDA as an executable extension of the benchmark, connecting multidimensional evidence with objective-guided strategy selection and standardized implementation. Rather than providing a static ranking, xConDA supports benchmark-guided selection according to study design and analytical objective, followed by standardized preprocessing, covariate-adjusted DAA and, where applicable, multi-cohort integration. This connection between benchmarking and execution provides a practical route for translating methodological evidence into routine microbiome analysis. As a complement to our previous xMarkerFinder [40] framework for downstream cross-cohort biomarker modeling, validation, and interpretation, xConDA addresses the upstream challenge of selecting and implementing an appropriate DAA strategy under confounding and multi-cohort settings.

Several limitations define the scope of these recommendations. Although our simulations were informed by empirical microbiome data and incorporated diverse confounding structures, they cannot capture all data-generating mechanisms, including unmeasured confounding, nonlinear effects and complex ecological dependencies [36, 37]. The empirical reference evaluations provide complementary biological evidence but do not constitute absolute ground truth. Our framework also addresses computational adjustment of measured covariates and cannot substitute for careful study design or comprehensive metadata collection. Moreover, the current benchmark primarily evaluates relative taxonomic abundance profiles. Quantitative microbiome profiling can directly capture microbial-load variation and address compositional biases that cannot be fully resolved computationally [1, 3, 22], making extension to quantitative data an important future direction. Method inclusion and preprocessing configurations were necessarily finite: DAA methods were restricted to those published by January 2025, and preprocessing was limited to widely used non-rarefaction approaches [16]. For multi-cohort analyses, our simulations emphasized pronounced between-cohort heterogeneity while assuming a common direction of phenotype-associated effects; settings with weak cohort effect, including potentially discordant phenotype effects across cohorts, remain to be evaluated as data-processing workflows improve. Longitudinal and repeated-measure designs, heterogeneous cohort effects, functional profiles and strain-level variation will therefore require dedicated evaluation. Within these boundaries, our results support treating confounder adjustment, preprocessing, statistical modelling and cross-cohort integration as components of a complete analytical strategy and using multidimensional benchmark evidence to guide more transparent and reproducible microbiome DAA.

## Methods

### 1 Benchmark design and construction of single-cohort DAA strategies

We benchmarked confounder-aware differential abundance analysis at the strategy level, defining each single-cohort DAA strategy as a prespecified combination of an input or preprocessing procedure and a statistical DAA model. Six benchmark-defined input schemes were considered: unnormalized count input (count), centered log-ratio (CLR) transform; cumulative sum scaling (CSS) [41]; log2 transformation (LOG); trimmed mean of M-values (TMM) [42]; and total sum scaling (TSS). The exact mathematical operations, pseudocounts, scaling conventions, and treatment of method-specific internal normalization for each input scheme are provided in Supplementary Table 1. DAA methods were eligible for inclusion if they had been published and were available by January 2025, supported explicit covariate adjustment, and were computationally feasible within the benchmark pipeline. Fourteen eligible DAA methods were initially considered in combination with the six input schemes. DACOMP [43] and mixMC [44] were excluded because the implementations considered did not provide the required multivariable covariate-adjusted feature-level testing framework. Songbird [45] supports covariate-aware multinomial regression but was excluded because its differential-ranking output does not provide the conventional feature-level significance calls required for the common FDR-based benchmark. ANCOM [46] was excluded because of practical limitations in significance output and computational efficiency. Because not all methods were compatible with all input types, combinations that violated method-specific statistical assumptions or software requirements, or that showed recurrent computational failure under the benchmark settings, were excluded. In total, 71 feasible strategies were retained for evaluation (Table 1). The complete set of strategies and their standardized labels is provided in Supplementary Table 1.

Within each dataset, all strategies were evaluated using the same phenotype definition, covariate set, feature-filtering rule, and multiple-testing threshold. Strategy-specific differences were therefore restricted to the input or preprocessing procedure and the statistical DAA model. Retained strategies were classified into two input-configuration types: default input configurations following developer-recommended or documented workflows and extended input configurations combining external preprocessing procedures with downstream DAA models. Extended input configurations were retained only when computationally feasible and compatible with software requirements, and should be interpreted as empirically evaluated extensions rather than standard workflows endorsed by the original method developers.

The following subsections describe the modeling framework, input requirements, how confounders were incorporated, internal normalization, non-default parameters, and the external preprocessing schemes retained in the final strategy combinations. The 14 methods were grouped according to their methodological origin and typical application context: (1) microbiome-tailored methods, comprising ALDEx2 [12], ANCOM-BC2 [14], Corncob [13], fastANCOM [47], MaAsLin2 [48], VTwins [49], ZicoSeq [15]; (2) RNA-seq-derived methods commonly adapted for microbiome differential abundance analysis (DAA), comprising limma [50], edgeR [42], DESeq2 [51]; and (3) classical statistical methods commonly used in microbiome studies, comprising linear fixed-effects model (LFEM), linear model with interaction terms (LM-inter), linear mixed-effects model (LMEM), stratified Wilcoxon test.

#### 1.1 Microbiome-tailored methods

##### ALDEx2

ALDEx2 [12] accounts for compositionality by generating Monte Carlo instances from a Dirichlet posterior distribution, thereby propagating within-sample uncertainty in count-derived relative abundances into hypothesis testing, while between-sample variation is estimated from biological replicates. The Monte Carlo–sampled relative abundances are then transformed using centered log-ratio transformation. Covariate adjustment was performed using the *aldex.glm* function, with covariates included in the model formula and default parameters otherwise retained. Because internal Dirichlet Monte Carlo sampling and CLR transformation are conducted, ALDEx2 was applied only to count input. Accordingly, ALDEx2_count was classified as the standard configuration, yielding one ALDEx2 strategy.

##### ANCOM-BC2

ANCOM-BC2 [14] estimates taxon-specific log-fold changes from compositional sequencing data using abundance estimates corrected for sampling-fraction and taxon-specific sequencing-efficiency biases, with variance regularization applied to stabilize statistical inference. Analyses were performed using the *ancombc2* function from the ANCOMBC package, with *lib_cut* set to 100 to exclude samples with total library sizes below this threshold. For confounder adjustment, the fix_formula parameter included the primary group variable and relevant covariates as fixed effects. Count input was classified as the default configuration, consistent with the method’s native count-based workflow, whereas CSS- and TMM-preprocessed inputs were classified as extended configurations, yielding three ANCOM-BC2 strategies.

##### Corncob

Corncob [13] models taxon-specific proportions using taxon-wise beta-binomial regression, with separate mean and dispersion parameters estimated by maximum likelihood. Differential abundance was assessed using the *bbdml* function by testing the group effect in the mean model, with confounders included in both the mean model (*formula*) and dispersion model (*phi.formula*) to account for covariate-associated shifts in taxon proportions and extra-binomial variation. Because Corncob models taxon counts relative to sample totals, it was evaluated using count input only. Accordingly, Corncob_count was classified as the default input configuration, yielding one Corncob strategy.

##### fastANCOM

fastANCOM [47] provides a computationally efficient implementation of ANCOM that preserves its log-ratio-based rationale while approximating its inference through taxon-wise models for log-transformed counts instead of exhaustive pairwise log-ratio model fitting. Analysis was performed using the *fastANCOM* function with default parameters. Confounder adjustment was performed through the *Z* parameter, which included covariates in the regression model. Count input was classified as the default input configuration, consistent with the method’s native count-based workflow, whereas CSS-, TMM-, and TSS-preprocessed inputs were classified as extended input configurations, yielding four fastANCOM strategies.

##### MaAsLin2

MaAsLin2 [48] offers a regression framework for microbiome multivariable association testing, supporting multiple statistical models, including linear model (LM), compound Poisson linear model (CPLM), negative binomial model (NEGBIN), and zero-inflated negative binomial model (ZINB). MaAsLin2_GLM denotes the LM option implemented in MaAsLin2 using *glm* with *family = gaussian*. For each MaAsLin2 model option, the phenotype and measured covariates were included in the *fixed_effects* argument for confounder adjustment, with the model type specified through the *analysis_method* argument in the *Maaslin2* function. Unless otherwise specified, other parameters were kept at their default values. Data preprocessing was performed using MaAsLin2’s internal *normalization* and *transform* arguments, without applying external preprocessing. The package-default workflow, consisting of GLM with TSS normalization and log transformation, was designated as the default input configuration. The remaining evaluated combinations were selected from the model, normalization, and transformation options natively implemented and permitted by MaAsLin2 and were therefore included as extended input configurations. In total, eighteen feasible MaAsLin2 strategies were evaluated (Table 1, Supplementary Table 1).

##### VTwins

VTwins [49] constructs within-cohort “virtual twin” sample pairs with similar microbial profiles but different phenotypes using K-nearest neighbor (KNN)-based matching to reduce sample-level heterogeneity, followed by permutation-based paired differential abundance testing. This process was implemented using the *pair_find* function from the VTwins package, with *ShuffleTime* set to 1000 and other parameters kept at their default values. Because the KNN-based matching procedure operates directly on the supplied abundance matrix, its results may depend on the input scale. VTwins specifies relative-abundance input and illustrates total-sum normalization. Accordingly, TSS input was classified as the default configuration, whereas count, CLR, LOG, and TMM inputs were evaluated as extended configurations, yielding five VTwins strategies.

##### ZicoSeq

ZicoSeq [15] performs differential abundance testing using an empirical Bayesian log-ratio regression framework for compositional microbiome data, in which reference taxa are selected by minimizing residual variance and microbial abundance ratios are estimated using a Beta mixture prior with posterior sampling. The phenotype and measured covariates were included in the regression design for confounder adjustment, and software-reported permutation-based FDR-adjusted *P* values were used. We applied the *ZicoSeq* function with default parameters, except that a square-root transformation was specified using *link.func = list(function(x) x^0.5)*. The *feature.dat.type* parameter was specified according to the type of input data. Count input was classified as the default configuration, consistent with ZicoSeq’s native count-based workflow, whereas CSS-, TMM-, and TSS-preprocessed inputs were classified as extended configurations, yielding four ZicoSeq strategies.

#### 1.2 RNA-seq-derived methods

##### limma

limma [50] performs differential abundance testing using empirical Bayes–moderated linear models. The design matrix included the phenotype and measured covariates for confounder adjustment. Models were fitted using the *lmFit* function, followed by empirical Bayes moderation with *eBayes*, and differential abundance was assessed by testing the phenotype coefficient. Adjusted P values were extracted using the *topTable* function with BH correction. Because limma does not prescribe a microbiome-specific choice among the evaluated input schemes, all six were classified as extended input configurations, yielding six limma strategies.

##### edgeR

edgeR [42] performs differential abundance testing using negative binomial generalized linear models with quasi-likelihood (QL) inference. The design matrix included the phenotype and measured covariates for confounder adjustment. Dispersion was estimated using the *estimateDisp* function, the model was fitted using *glmQLFit*, and differential abundance was assessed using QL F-tests through *glmQLFTest*. TMM-preprocessed input was classified as the default configuration, consistent with edgeR’s recommended normalization approach, whereas count-, CSS-, and TSS-based inputs were classified as extended configurations, yielding four edgeR strategies.

##### DESeq2

DESeq2 [51] models count data using negative binomial generalized linear models, with median-of-ratios size factors correcting library-size differences and mean– dispersion trend shrinkage stabilizing dispersion estimation. The design formula included the phenotype and measured covariates for confounder adjustment. Differential abundance was assessed using the Wald test. This workflow was implemented using the *DESeq* function from the DESeq2 Bioconductor package with default parameters. Because DESeq2 is designed for raw count input and performs size-factor normalization internally, only the DESeq2_count strategy was evaluated and was classified as the default input configuration.

#### 1.3 Classical statistical methods

##### Linear fixed effect model (LFEM)

LFEM evaluated differential abundance by fitting ordinary least squares (OLS) linear models for each feature, with the phenotype and measured covariates included as fixed effects for confounder adjustment. Model fitting was performed using the *lm* function from base R with the formula feature ∼ label + confounder. As LFEM does not prescribe a microbiome-specific preprocessing scheme, all six input schemes were classified as extended input configurations, yielding six LFEM strategies.

##### Linear mixed effect model (LMEM)

Linear mixed-effects models were fitted for each feature, with the phenotype and covariates specified in the model formula; continuous covariates were treated as fixed effects and applicable categorical covariates as random intercepts to account for grouped observations. Models were fitted using the *lmer* function from the *lmerTest* package with the formula *feature ∼ label + continuous covariates + (1 | categorical covariate)*. As LMEM does not prescribe a microbiome-specific preprocessing scheme, all six input schemes were classified as extended input configurations, yielding six LMEM strategies.

##### Linear model with interaction term (LM-inter)

LM-Inter extends LFEM by adding phenotype–covariate interaction terms to model potential effect modification. For each taxon, a linear model was fitted using the *lm* function from base R with the formula *feature ∼ label + covariates + label:covariates*. The phenotype association was assessed using the BH-adjusted *P* value of the phenotype main-effect coefficient. As LM-Inter does not prescribe a microbiome-specific preprocessing scheme, all six input schemes were classified as extended input configurations, yielding six LM-Inter strategies.

##### Stratified Wilcoxon test

The stratified Wilcoxon test is a non-parametric method that accounts for confounding. Continuous confounders were discretized and combined with categorical confounders into a single stratification factor. For each taxon, stratified Wilcoxon testing was performed using the *wilcox_test* function from the coin package with the formula *feature ∼ label | confounder_block*, where *confounder_block* denotes the combined stratification factor derived from all confounders. As this test does not prescribe a microbiome-specific preprocessing scheme, all six input schemes were classified as extended input configurations, yielding six strategies.

### 2 Construction and implementation of multi-cohort DAA strategies

We extended the single-cohort benchmark to multi-cohort settings by defining each multi-cohort DAA strategy as a combination of three components: an input or preprocessing procedure, a statistical DAA model, and a cross-cohort integration approach. The 12 benchmark-prioritized single-cohort strategies were carried forward and combined with two major classes of multi-dataset integration frameworks: mega-analysis, which pools individual-level data across cohorts and explicitly models batch effects, and meta-analysis, which aggregates study-level association statistics. This design yielded 63 multi-cohort DAA strategies.

#### 2.1 Mega-analysis strategy (sample-level integration)

For mega-analysis, feature-aligned sample-level abundance profiles from all cohorts were pooled into a unified matrix, and each benchmark-prioritized single-cohort strategy was applied with cohort identity included as a fixed-effect covariate alongside harmonized confounders. This pooled framework accounts for cohort-specific baseline differences while leveraging all samples in a single model. Mega-analysis strategies were denoted by the suffix _Mega.

#### 2.2 Meta-analysis strategies (study-level integration)

For meta-analysis, each DAA strategy was first applied independently within each cohort using the same phenotype definition and harmonized covariate specification. Cohort-specific P values, effect estimates and standard errors, where available, were then combined across cohorts. We evaluated two broad types of meta-analysis: P-value-based integration and effect-size-based integration. These were implemented in R packages metap (v1.11) and metafor (v4.8-0):

##### P-value-based meta-analysis

∎ **Stouffer’s method** combines Z-transformed P values across studies, with weights proportional to the square root of each study’s sample size, allowing larger cohorts to contribute proportionally more information.
∎ **Fisher’s method** aggregates evidence through the sum of -2log(p), providing a sensitive test for detecting signals present in multiple cohorts.

Fisher’s and Stouffer’s methods combine two-sided P values without encoding effect direction. Their results should therefore be interpreted alongside cohort-specific effect estimates.

##### Effect-size–based methods

∎ **Empirical Bayes (EB)** integrates effect sizes under an empirical prior, yielding shrinkage-based estimates robust to study-level variability.
∎ **Restricted maximum likelihood (REML)** estimates between-study variance by maximizing the restricted likelihood, providing stable inference in settings with heterogeneous effect size.
∎ **Paule–Mandel (PM)** iteratively matches observed and expected effect-size variability, delivering reliable variance estimates even under substantial heterogeneity.

These five frameworks cover common meta-analysis approaches with different assumptions about heterogeneity modeling, evidence pooling, and study weighting.

### 2.3 Construction of end-to-end multi-cohort DAA pipeline

To construct multi-cohort DAA pipelines, we paired the 12 benchmark-prioritized single-cohort strategies with compatible cross-cohort integration approaches. All 12 benchmark-prioritized single-cohort DAA strategies were compatible with mega-analysis, yielding 12 mega-analysis strategies, and with Fisher and Stouffer P-value integration, yielding 24 P-value–based meta-analysis strategies. Effect-size–based meta-analysis was restricted to strategies that reported comparable phenotype-effect estimates and standard errors; nine eligible strategies, including ANCOM-BC2, DESeq2, and MaAsLin2 variants, were paired with three random-effects estimators, REML, PM, and EB, yielding 27 effect-size–based meta-analysis strategies. In total, 63 multi-cohort DAA strategies were constructed and named by appending an integration suffix to the corresponding single-cohort strategy: _Mega for mega-analysis and _Meta-F, _Meta-S, _Meta-R, _Meta-P, and _Meta-E for Fisher, Stouffer, REML, PM, and EB integration, respectively. The full strategy list is provided in Supplementary Table 14.

### 3 Benchmarking datasets and simulation design

#### 3.1 Real-data-informed simulation framework

We used SparseDOSSA2 [36] to generate synthetic microbiome count profiles for benchmarking. SparseDOSSA2 estimates generative parameters from empirical microbiome data and produces synthetic count matrices that preserve key microbial community characteristics, including sparsity, compositional structure, abundance distributions, and feature–feature correlations. This generative framework enabled the construction of simulation scenarios with systematically varied sample sizes, confounding structures, and effect settings. We constructed dedicated simulation settings to evaluate single-cohort accuracy, perturbation-based stability, and multi-cohort accuracy. Complete parameter definitions and scenario configurations are provided in Supplementary Tables 2 and 9.

##### Common simulation settings

The simulations were parameterized using gut microbiome community structures derived from real datasets, providing high-diversity microbial profiles representative of human gut metagenomic studies. Each base simulation generated a synthetic population of 10,000 samples with 1,000 microbial features and a median sequencing depth of approximately 3×10^6^ reads. Scenario-specific datasets were randomly subsampled from the corresponding synthetic population to obtain the required sample sizes while retaining sequencing-depth variability and introducing finite-sample sampling noise.

In the primary full-factorial simulations, phenotype-associated effects were introduced into 20% of microbial features to create a signal-rich setting with substantial compositional perturbation. Because perturbation of multiple features can induce secondary relative-abundance shifts among unperturbed features, this design provided a challenging setting for evaluating false-positive control. A complementary sensitivity analysis reduced the proportion of phenotype-associated features to 5% to evaluate strategy performance under a substantially sparser signal setting, as described below. The 5% setting represented a distinct sparse-signal challenge, requiring methods to detect limited true signals while maintaining adequate error control. Both enriched and depleted features were generated, with asymmetric bidirectional effects used to reflect unequal effect distributions commonly observed in empirical microbiome studies. Comparison of the two settings was used to determine whether the main recommendations remained consistent across different proportions of differentially abundant features.

Continuous and categorical confounders were generated to be associated with both phenotype status and microbial abundances. Phenotype effects and confounder effects were parameterized separately but were allowed to act on overlapping feature subsets. Features directly assigned a phenotype effect constituted the ground-truth DA set, whereas features affected only by confounders represented structured nuisance signals for evaluating covariate adjustment.

##### Configurations of single-cohort accuracy evaluation

To evaluate performance under diverse single-cohort conditions, we varied five factors chosen to cover key aspects of study design, confounding structure, and disease-context heterogeneity: sample size, case–control ratio, confounder complexity, confounding strength, and real-data-derived disease-context template. Sample size was varied from small to well-powered designs, and case–control ratios ranged from balanced to strongly imbalanced group designs. Confounder complexity was varied by changing the number and type of confounders, whereas confounding strength ranged from marginal to dominant relative to the phenotype-associated effect, thereby increasing the difficulty of confounder adjustment.

Five disease-context templates were used to represent microbiome-associated disease contexts with distinct empirical microbiome community structures: autism spectrum disorder (ASD; neurodevelopmental), colorectal cancer (CRC; oncological), inflammatory bowel disease (IBD; gastrointestinal inflammation), rheumatoid arthritis (RA; autoimmune), and type 2 diabetes (T2D; metabolic) [52]. These labels denote the real-data parameterizations used to define the background microbial distributions and should not be interpreted as simulations of complete disease-specific biological mechanisms.

The full-factorial design across the five factors yielded 250 simulation scenarios. Each scenario was repeated ten times using independent random seeds, with performance metrics calculated separately for each replicate before aggregation. Exact factor levels and parameter combinations are provided in Supplementary Table 2.

##### Sparse-signal sensitivity analysis

To assess whether strategy prioritization depended on the prespecified proportion of differentially abundant features, we conducted a complementary sensitivity analysis in which this proportion was reduced from 20% to 5%. Six representative scenarios were constructed by combining three sample designs—small balanced, small imbalanced, and large balanced—with low or high confounding burdens, defined by both the number and effect strength of confounders. These scenarios captured key challenges to sparse-signal detection arising from limited sample size, group imbalance, and confounding. Each scenario was repeated ten times using independent random seeds, while the phenotype-associated effect size and other data-generation procedures were kept consistent with the primary simulations. Corresponding parameter configurations are reported in Supplementary Table 2.

##### Configurations of single-cohort stability evaluation

We assessed strategy stability using one-factor-at-a-time perturbation analyses. For each analysis, one factor was varied across ordered levels while all other parameters were fixed at a common baseline comprising 150 cases and 150 controls, two continuous and one categorical confounder, a moderate phenotype effect and a weaker confounding effect. The most challenging level of each factor was additionally used for worst-case performance evaluation.

Per-group sample size was varied across 25, 50, 100, 150, 200, 300, and 500 samples. Case–control imbalance was evaluated under a fixed total sample size, ranging from balanced allocation to 30 cases and 570 controls. Phenotype-effect magnitude was varied from 2 to 8, whereas confounder-effect magnitude was varied from 1 to 5, according to the simulation scale defined in Supplementary Table 9. Confounder complexity was increased stepwise by including one continuous confounder together with one to four categorical confounders.

In addition, performance was evaluated across feature-prevalence strata without applying the standard low-prevalence filter. Features were grouped by observed prevalence into intervals of 0–0.05, 0.05–0.10, 0.10–0.20, and 0.20–1.00. Because prevalence was not directly manipulated as a simulation factor, this analysis evaluated method performance across feature groups stratified by observed prevalence.

Full parameter configurations are reported in Supplementary Table 9.

##### Multi-cohort simulation

To evaluate multi-cohort DAA under pronounced cohort heterogeneity, we generated a focused stress-test setting comprising 1,000 samples distributed across five cohorts. Cohort-specific effects were represented by indicator variables that induced abundance shifts in predefined feature subsets. The affected subsets were allowed to overlap partially or remain cohort-specific, thereby generating feature-dependent cohort bias rather than a uniform global batch shift.

Cohort effects were specified to vary in magnitude across cohorts and to be stronger overall than the common phenotype effect, reflecting settings in which heterogeneous between-study variation dominates the biological association of interest. To retain an unambiguous benchmark ground truth, the phenotype effect was specified as common across cohorts; scenarios in which the phenotype effect was absent in some cohorts or reversed in direction were not simulated. Additional measured confounders had weaker effects than the phenotype and were restricted to three covariates shared across all cohorts: two continuous and one categorical variable. This design reproduced two common challenges in cross-study microbiome analysis: strong cohort-level heterogeneity and limited availability of harmonizable covariates. The common phenotype effect and measured-confounder effects followed the fixed settings used in the perturbation-stability simulations, and samples were randomly partitioned across five cohorts to define cohort sizes and case–control allocations.

#### 3.2 Real-world benchmark datasets

The real-data benchmark comprised two complementary dataset collections. Reference datasets with established or curated microbial signatures were used to evaluate biological signal recovery, whereas datasets without predefined reference signatures were used to assess within-dataset consistency and reproducibility under cohort heterogeneity.

##### Reference datasets with established biological signatures

Six datasets were included in the reference-based evaluation. These comprised two 16S rRNA gene sequencing datasets, including one vaginal microbiome dataset comparing healthy vaginal communities with bacterial vaginosis [53] and one paired oral microbiome dataset comparing supragingival and subgingival dental plaque from the same participants [54], as well as four independent shotgun metagenomic datasets comparing individuals with CD with healthy controls (PRJNA385949, PRJEB1220, PRJNA398089, and PRJEB18780) [9, 55–58].

The vaginal and oral datasets provided established clinical or ecological reference patterns, whereas the four CD datasets were used to evaluate recovery of independently curated disease-associated taxa representing concordant microbial shifts reported across multiple studies. The CD reference set was derived exclusively from external studies that did not overlap with the four evaluation cohorts, as described in Section 5.2. Datasets were obtained from publicly available resources using the R packages *curatedMetagenomicData* (v3.10.0) [31], *HMP16SData*[54], and *MicrobiomeBenchmarkData* [59].

##### Datasets for consistency evaluation

Within-dataset consistency and multi-cohort integration reproducibility were evaluated using 36 human gut metagenomic disease-control datasets obtained from curatedMetagenomicData (v3.10.0) [31]. Species-level taxonomic profiles in curatedMetagenomicData were generated using MetaPhlAn3 [60]. After applying the criteria, 36 eligible cohorts were retained. Because several cohorts contained multiple disease phenotypes, each eligible disease-control contrast within a cohort was treated as a separate disease-control dataset for the within-dataset consistency analysis. This procedure yielded 50 disease-control datasets derived from 36 human gut metagenomic cohorts. Phenotype definitions and available covariates were harmonized before analysis. Additionally, the MetaPhlAn3 taxon counts provided by curatedMetagenomicData were generated by multiplying relative abundances by the corresponding sample-level sequencing depths [31].

For the multi-cohort integration consistency analysis, we retained disease indications represented by at least four independent cohorts: adenoma (five cohorts), CRC (11 cohorts), IBD (five cohorts) and T2D (four cohorts). These cohorts were used to evaluate reproducibility by applying the same multi-cohort DAA strategy to independently assembled groups of cohorts.

After accounting for overlap between the four Crohn’s disease independent cohorts and the curatedMetagenomicData collection, the benchmark comprised 38 unique real-world datasets.

### 4 Simulation performance assessment

#### 4.1 Accuracy evaluation metrics

In each simulation scenario, features assigned a direct phenotype-associated effect were defined as ground-truth differentially abundant (DA) features, whereas all remaining features were classified as non-DA. Features affected by simulated confounders but not directly by the phenotype were therefore retained in the non-DA class, enabling evaluation of confounding-induced false discoveries. For each strategy, detected DA features were compared with the simulation-defined ground truth to calculate true positives (TP), false positives (FP), true negatives (TN), and false negatives (FN), from which threshold-dependent performance metrics were derived. Features with Benjamini–Hochberg-adjusted (BH-adjusted) *P*-values <0.05 were considered significant for threshold-based metrics. Each scenario was independently simulated ten times, with metrics calculated per replicate and averaged for performance estimation.

##### Area Under the Receiver Operating Characteristic Curve (AUROC)

AUROC was used to assess threshold-independent discrimination between true DA and non-DA features. For each strategy, features were ranked using 1 − the adjusted P values generated by its standard analysis workflow, with higher scores indicating stronger evidence for differential abundance. Taxa omitted because of method-specific filtering or model-fitting failure were retained in the common evaluation set and assigned an adjusted P value of 1. ROC curves were constructed by comparing these scores with the simulation-defined DA labels.

##### Area Under the Precision–Recall Curve (AUPR)

AUPR was calculated using the same ranking scores, with true DA features treated as the positive class, and was used to assess precision–recall performance under DA feature sparsity.

##### False Positive Rate (FPR)

FPR was defined as the proportion of non-DA features incorrectly identified as significant:

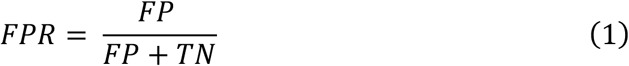

##### False Discovery Rate (FDR)

FDR was defined as the proportion of false positives among all detected DA features:

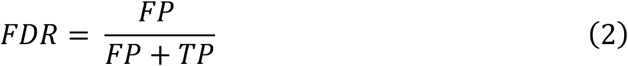

##### Macro-F1

Macro-F1 was used to summarize classification performance across DA and non-DA classes while reducing dominance by the majority class. The F1-score was calculated separately for the positive and negative classes:

For the differential (positive) class,

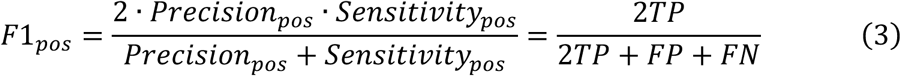

For the non-differential (negative) class (treating “non-differential” as the target class),

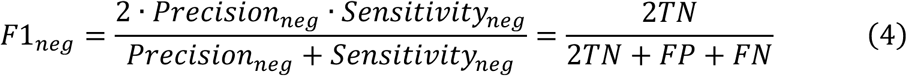

Then,

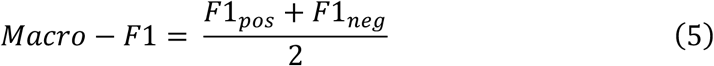

##### Matthews Correlation Coefficient (MCC)

MCC was used to evaluate agreement between detected and true DA labels while accounting for all four entries of the confusion matrix:

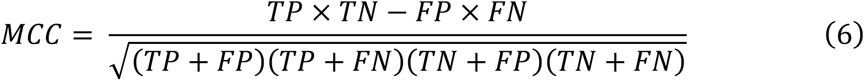

MCC ranges from −1 to 1, with higher values indicating better classification performance, and is particularly informative under class imbalance.

##### Sensitivity

Sensitivity, also known as the true positive rate, was defined as the proportion of truly DA features correctly detected

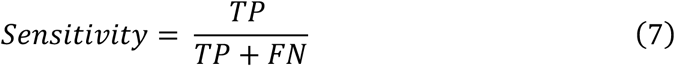

#### 4.2 Composite ranking of simulation accuracy

We used Technique for Order Preference by Similarity to Ideal Solution (TOPSIS) to integrate AUROC, AUPR, MCC, and Macro-F1 into a composite simulation-accuracy score. These summary accuracy metrics capture complementary aspects of performance: AUROC and AUPR quantify threshold-independent discrimination, whereas MCC and Macro-F1 assess balanced classification at the prespecified significance threshold.

For each simulation scenario, metric values were first averaged across the ten replicates, and a decision matrix was then constructed with strategies as rows and the four metrics as criteria. Metrics were min–max normalized across strategies within each scenario and assigned equal weights. The positive and negative ideal solutions were defined as the column-wise maximum and minimum of the weighted normalized matrix, respectively. For each strategy, the Euclidean distances to the positive and negative ideal solutions were denoted by *d*^+^ and *d*^−^, respectively. The TOPSIS score was defined as:

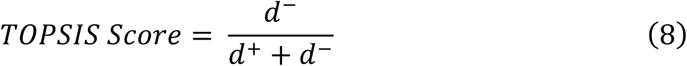

Strategies were ranked within each simulation setting by TOPSIS score, with higher scores indicating better performance. Overall strategy rankings were derived from the mean scenario-specific TOPSIS score across all simulation scenarios, with metric-level means and standard deviations reported as summary statistics.

### 4.3 Perturbation-based stability evaluation

We evaluated strategy stability under systematic perturbations of sample size, phenotype effect size, confounding effect size, confounder complexity, case–control ratio and feature prevalence, using the perturbation settings described in Section 3.1. For each factor, one parameter was varied across ordered levels while all other parameters were held at the common baseline. Each level was simulated ten times, and replicate-level metrics were averaged before trend analysis.

Performance across perturbation levels was summarized using the Area Under the Metric-Trend Curve (AUMTC) as:

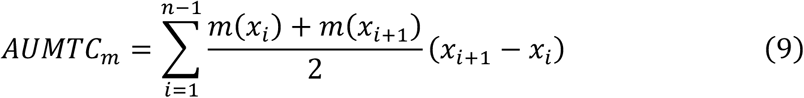

where *x_i_* denotes the i-th ordered factor level, n is the total number of levels, and *m*(*x_i_*) denotes the min-max-normalized metric value at *x_i_*. Ordered factor levels were assigned numeric values at equal intervals from 0 to 1 according to their ranks. AUMTC was computed for each metric *m*, including MCC, Macro-F1, AUPR, AUROC, sensitivity, FDR, and FPR. For FDR and FPR, normalized values were reversed so that larger values consistently indicated better performance.

AUMTC was calculated separately for AUROC, AUPR, MCC, Macro-F1, sensitivity, FPR, and empirical FDR. Because AUMTC reflects both overall performance and performance maintenance across perturbation levels, it was interpreted as a trend-integrated stability measure. AUMTC values for AUROC, AUPR, MCC, and Macro-F1 were further integrated using equal-weight TOPSIS to obtain a composite stability score, which was used to rank strategies under each perturbation factor.

Worst-case performance was additionally evaluated at the most challenging prespecified level of each perturbation factor. The tested levels were 25 samples per group, a case–control ratio of 30:570, a phenotype effect of 2 and a confounding effect of 5. The most complex covariate structure comprised one continuous and four categorical variables, and the lowest feature-prevalence stratum was <0.05 (Supplementary Table 12). Together, AUMTC and worst-case performance jointly characterized strategy stability by summarizing performance maintenance across gradual perturbations and at the most challenging prespecified settings, respectively.

### 5 Empirical benchmark performance evaluation

The 71 single-cohort DAA strategies were applied to empirical benchmark datasets for reference-based recovery and consistency evaluation. Because exact ground truth was unavailable, strategy performance was assessed using predefined biological reference signatures and reproducibility-based criteria, including reference-pattern recovery [19], reference taxa recovery, and consistency evaluation.

#### 5.1 Reference-pattern recovery assessment

Reference-pattern recovery was evaluated in real datasets for which expected microbial shifts were supported by established ecological, culture-based, or clinical evidence. For each dataset, we defined direction-specific reference taxa and assessed whether each strategy recovered these taxa in the expected direction. Performance was summarized using the recovery breadth, defined as the number of reference taxa recovered in the expected direction, and enrichment significance.

For each dataset, strategy and reference direction, taxa detected as differentially abundant in the expected direction were tested for enrichment of the corresponding reference taxa. All taxa tested by the strategy in that dataset were used as the background. For this test, taxa in the background were cross-classified by reference-set membership and detection in the expected direction, yielding a 2 × 2 contingency table with four entries: reference taxa recovered in the expected direction (a), non-reference DA taxa detected in the expected direction (b), reference taxa not recovered in the expected direction (c), and non-reference taxa not detected as DA in the expected direction (d). Enrichment significance was assessed using one-sided Fisher’s exact tests with the alternative hypothesis set to greater. P values were adjusted across strategies within each dataset and reference direction using the Benjamini–Hochberg (BH) procedure.

To jointly account for recovery breadth and enrichment significance, we defined the Reference Recovery Enrichment Score (RRES). Direction-specific recall was calculated as:

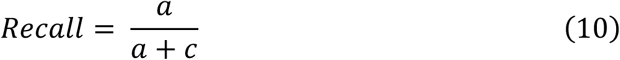

where (*a* + *c*) denotes the number of reference taxa for that direction. The direction-specific RRES was defined as:

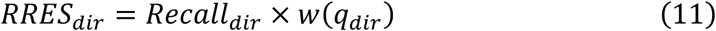

where *q_dir_* is the BH-adjusted Fisher’s exact test P value for that direction, and *w*(*q_dir_*) maps the enrichment evidence to a bounded weight in [0, 1]:

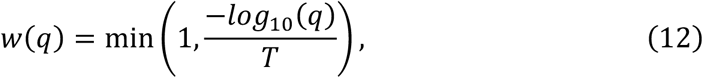

Here, smaller q values yield larger weights, with the saturation parameter set to T = 5, so that weights are capped at 1 for q ≤ 10⁻⁵ and extremely small q values do not further dominate the score.

For datasets with bidirectional reference patterns, direction-specific scores were combined using the geometric mean:

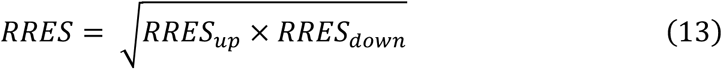

Strategies were ranked by RRES to prioritize methods with higher recovery breadth in the expected direction and stronger enrichment evidence.

**(1) Oral microbiome: supragingival vs. subgingival plaque**

Reference taxa were annotated by oxygen requirement as aerobic, anaerobic, facultative anaerobic, or unassigned using the nychanesmicrobiome resource [61] (https://github.com/waldronlab/nychanesmicrobiome). Aerobic taxa were used as reference taxa for supragingival enrichment, and anaerobic taxa were used as reference taxa for subgingival enrichment. Facultative anaerobic and unassigned taxa were retained in the tested background but were not used as direction-specific reference taxa. For each DAA strategy, we calculated recovery breadth, enrichment significance, and RRES for each direction, and then summarized them as a combined bidirectional score.

This dataset consisted of paired supragingival and subgingival plaque samples from the same individuals and was analyzed as a covariate-free reference-pattern evaluation. In the absence of additional confounders, strategy variants differing only in confounder-adjustment structure, including mixed-effects, fixed-effects, and interaction-term formulations, reduced to equivalent model specifications. These equivalent variants were collapsed before summarization, yielding 59 non-redundant strategies for evaluating recovery of the expected ecological pattern.

**(2) Vaginal microbiomes: healthy vaginal vs. bacterial vaginosis**

Reference taxa were defined using curated BV- and HV-associated taxa from MicrobiomeBenchmarkData [59], which integrates evidence from culture-based studies, microscopy, and previous large-scale 16S rRNA and metagenomic studies [62–67]. HV-associated taxa were used as reference taxa for HV enrichment, and BV-associated taxa were used as reference taxa for BV enrichment. For each DAA strategy, recovery breadth, enrichment significance, and RRES were calculated separately for the upward and downward directions and then summarized as a combined bidirectional score.

#### 5.2 Reference taxa recovery using a Crohn’s disease evidence compendium

CD was used to evaluate whether DAA strategies could recover reproducible disease-associated taxa in real metagenomic datasets. Because no exact taxon-level ground truth exists for CD, we constructed a compact positive reference set from external evidence rather than treating any single dataset as a gold standard. We synthesized evidence from 42 external CD studies, which collectively reported 387 differentially abundant species. Species names were harmonized to a common taxonomy, and only studies independent of the four evaluation datasets were used for reference curation to avoid information leakage. Species were retained as reference taxa if they were reported in at least three independent studies with concordant disease-associated direction. The resulting reference set was further supported by independent functional or mechanistic evidence relevant to CD [68–74]. The final reference set comprised four species: *Ruminococcus gnavus* and *Escherichia coli,* which were enriched in CD, *Bifidobacterium adolescentis* and *Faecalibacterium prausnitzii,* which were depleted in CD.

Stable recovery of these reference taxa was evaluated in four independent CD shotgun metagenomic datasets that were not used for reference curation. For each strategy and reference taxon, recovery was scored as 1 if the taxon was detected as differentially abundant in the expected direction in at least three of the four evaluation datasets, and 0 otherwise. The Stable Recovery Score (SRS) was calculated as the sum of these binary recovery scores across the four reference taxa:

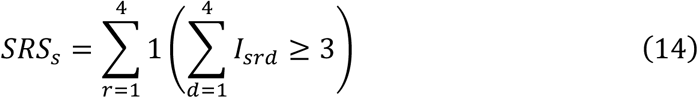

where *I_srd_* = 1 if reference taxon *r* was detected by strategy *s* as differentially abundant in the expected direction in evaluation dataset *d*, and *I_srd_* = 0 otherwise. Thus, SRS ranges from 0 to 4, with higher values indicating more consistent recovery of established CD-associated signals across independent cohorts. SRS was used as a positive-reference recovery assessment.

#### 5.3 Single-cohort within-dataset consistency

Single-cohort within-dataset consistency was evaluated for the 60 DAA strategies to assess reproducibility within individual studies under sample-level heterogeneity. Each dataset was randomly split into two equal, non-overlapping subsets across ten independent replicates. The two subsets from the same dataset were analyzed independently using the same single-cohort strategy, and significant taxa were defined as those with BH-adjusted *P* values < 0.05.

For each replicate, consistency between two detected DA sets identified from two independent subsets, A and B, was quantified using the Jaccard index. Each set comprised significant taxon–direction pairs, such that a taxon was considered overlapping only when it was detected in both subsets with the same direction of association:

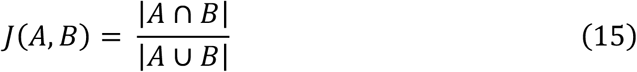

When both sets were empty, the Jaccard index was set to 0 to avoid rewarding strategies that appeared consistent only because they detected no DA taxa. The number of detected DA taxa was therefore recorded alongside the Jaccard index to distinguish reproducibility from discovery yield. Replicate-level Jaccard indices and DA counts were first summarized within each dataset and then aggregated across datasets using both mean and median values.

#### 5.4 Cross-cohort consistency of multi-cohort integration strategies

Cross-cohort consistency of multi-cohort DAA strategies was evaluated at two complementary levels. First, DA taxon detection consistency was assessed for all 63 multi-cohort DAA strategies to enable a common comparison across integration frameworks. Unlike single-cohort within-dataset consistency, which evaluates stability across sample-level resampling within the same study, this analysis assessed consistency across alternative combinations of independent cohorts using a cohort-level split-half resampling framework. For each disease represented by at least four cohorts, cohorts were randomly partitioned into two non-overlapping groups as evenly as possible across ten replicates. Each cohort group was analyzed separately using the same multi-cohort DAA strategy, including its corresponding preprocessing procedure, DAA model, and cross-cohort integration approach. For each split, significant taxa were defined as those with BH-adjusted *P* < 0.05. The Jaccard index between the significant taxa identified from the two cohort groups was calculated based on taxon identity, and the number of detected DA taxa was recorded. These measurements quantified the consistency of DA taxon detection across different combinations of independent cohorts.

Second, we conducted a direction-concordant detection-consistency analysis as a sensitivity analysis for the 39 strategies that provided signed integrated association estimates. A taxon was considered concordantly identified only when it was significant in both cohort groups and showed the same direction of association. A direction-aware Jaccard index was then calculated to evaluate the reproducibility of both taxon selection and association direction across cohort combinations. For both analyses, values were first averaged across cohort splits within each disease and then averaged across diseases.

### 6 Calculation of overall performance scores

To summarize multidimensional benchmark performance, an overall score was calculated separately for single-cohort and multi-cohort strategies using TOPSIS. All component scores were min–max normalized across strategies before integration. For single-cohort strategies, the overall score integrated the simulation accuracy TOPSIS score, RRES scores from the oral and vaginal reference benchmarks, the SRS from the Crohn’s disease reference benchmark, the mean Jaccard index for within-dataset reproducibility, and six AUMTC-based stability scores corresponding to sample size, case–control ratio, phenotype effect size, confounding effect size, confounder complexity, and feature prevalence. For multi-cohort strategies, the overall score integrated the simulation accuracy TOPSIS score and the mean Jaccard index for cross-cohort consistency. Equal weighting was applied at the level of individual component criteria, with each score treated as a distinct and complementary source of evidence without preferentially emphasizing any performance aspect, benchmark, or perturbation factor. Higher overall scores indicated performance profiles closer to the ideal solution. The overall score was used to summarize multidimensional performance and was not treated as the sole basis for strategy recommendation.

### 7 Sibling-informed reference analysis for the active CD case study

Following the integrated single-cohort benchmark, we performed an active Crohn’s disease (CD) case study using data from our previously published sibling-matched cohort [8]. We used two reference sets previously defined in that study. Sibling-matched reference taxa comprised taxa identified as differentially abundant between patients with active CD and their unaffected siblings and were used as putative CD-associated signals under a family-matched comparison. Family-design-derived confounding-sensitive reference taxa were defined as taxa differentially abundant between patients with active CD and unrelated controls but no longer differentially abundant in the sibling-matched comparison, representing putative familial or environmental signals.

Six benchmark-prioritized strategies spanning the general-purpose, high-confidence inference and broader signal-recovery branches of the decision framework—ANCOM-BC2_count, ANCOM-BC2_TMM, ANCOM-BC2_CSS, edgeR_CSS, MaAsLin2_CPLM_TSS and edgeR_count, were applied to the active CD versus unrelated-control comparison using their corresponding preprocessing and covariate-adjusted DAA procedures. DA taxa were defined at BH-adjusted *P* < 0.1. Recovery of sibling-matched reference taxa required both taxon identity and concordant association direction. For each strategy, we recorded the total number of detected DA taxa and the number of recovered sibling-matched reference taxa, and calculated the proportion of predefined confounding-sensitive taxa that remained non-differential. Analyses were repeated at BH-adjusted *P* < 0.2 as a sensitivity analysis.

### 8 Statistical analysis, visualization and reproducibility

Unless otherwise specified, microbial features detected in fewer than 10% of samples were removed before DAA, and this filtering rule was applied consistently across all strategies within each dataset. In the prevalence-stratified robustness analysis, features were retained and grouped by prevalence interval to assess strategy performance across different feature-prevalence levels.

For each DAA strategy, statistical significance was assessed at the feature level. Except for ZicoSeq, raw *P* values were adjusted across all tested features within each dataset, contrast and strategy using the BH procedure. For ZicoSeq, the software-provided permutation-based FDR-adjusted *P* values were used. For meta-analysis and mega-analysis strategies, BH correction was performed after cross-cohort integration across all tested features. In the rare instances where a strategy returned incomplete feature-level results or no valid output, the unavailable results were recorded as NA and treated as nonsignificant in all downstream evaluations. Features with adjusted *P* values below 0.05 were considered differentially abundant.

The impact of covariate adjustment on DAA performance was assessed using paired Wilcoxon signed-rank tests for each evaluation metric, comparing matched strategy-level results between unadjusted and covariate-adjusted analyses with statistical significance defined as *P* < 0.05.

All DAA strategies were executed within a unified benchmarking pipeline, with random seeds fixed for stochastic procedures where applicable. Analyses were performed using Python 3.8.18 and R 4.3.1. To support reproducibility, the complete analysis code is provided together with a Docker container documenting the computational environment, software dependencies and package versions used in this study.

Performance summaries and visualizations were generated using R packages including circlize v0.4.16, ComplexHeatmap v2.18.0, funkyheatmap v0.5.1, corrplot v0.95, ggplot2 v3.5.2 and fmsb v0.7.6. These packages were used to generate heatmaps, correlation heatmaps, bubble plots, radar charts and Circos-style plots.

## Data availability

All real datasets used in this study were obtained from publicly available resources, including the Bio-Med Big Data Center project OEP004514 and the R packages curatedMetagenomicData [31], HMP16SData [54], and MicrobiomeBenchmarkData [59]. Simulated datasets and the complete strategy-level differential abundance analysis outputs generated from all simulated and real datasets in this study are deposited in Zenodo (doi: 10.5281/zenodo.19883890). Processed performance metrics used for method evaluation are provided in the Supplementary Tables.

## Code availability

All analysis and visualization code, including Jupyter notebooks, Python scripts, R scripts and Docker configuration files, is publicly available at https://github.com/tjcadd2020/xConDA_benchmark.

## Acknowledgements

We thank the Tongji University Supercomputing Center and the Bioinformatics Supercomputing Center, School of Life Sciences and Technology, Tongji University, for providing computational resources and technical support.

## Author contributions

R.Z., N.J., G.P.Z. and G.Q.Z. conceived the study. X.Z. designed and performed the benchmark analyses, implemented the computational workflows, analyzed and interpreted the data, generated the figures and tables, and drafted the manuscript. X.Z. and B.Y. developed xConDA. L.L., Y.C., Y.S. and D.W. contributed to data curation, computational analyses, method implementation and validation. All authors critically revised the manuscript and reviewed and approved the final version.

## Ethics and Consent to Participate

This study used simulated datasets and previously published, publicly available microbiome datasets. No new human participants were recruited, and no new human samples were collected. Therefore, ethics approval and consent to participate were not applicable to the present study.

